# Temporally distinct CDX programmes preconfigure vagal and trunk neural crest

**DOI:** 10.64898/2026.07.13.738216

**Authors:** Irène Amblard, Christos M Kalaitzis, Irina Balaguer Balsells, Ivan Andrew, Ka Lok Choi, Hothri Ananyambica Moka, Laurence Game, Juan M Vaquerizas, Vicki Metzis

**Author notes:** equal contribution (Irène Amblard, Christos M Kalaitzis).

## Abstract

Neural crest cells (NCCs) are progenitor cells vital in establishing the head, heart, gut and peripheral nervous system of vertebrate embryos. Disruptions to NCC development underlie neurocristopathies, which constitute a wide array of congenital anomalies. Yet how NCCs acquire defined regional identities that enable them to generate distinct derivatives along the body axis remains unclear. Here, we identify an epiblast progenitor population in mouse embryos that transiently contributes to vagal neural crest cells and trunk-to-tail derivatives. Using single-cell spatial transcriptomics across successive stages of neural crest migration, we generate a cervicothoracic cell atlas that resolves vagal and trunk neural crest cells *in situ*. Combining *in vivo* lineage tracing with *in vitro* models of neural crest induction, we show that despite transiently sharing a lineage, vagal and trunk neural crest arise through separate mechanisms. Temporally discrete regionalisation events mediated by CDX transcription factors establish HOX states that define vagal versus trunk identity. These findings revise models of NCC formation by demonstrating that temporally separate epiblast regionalisation events preconfigure neural crest and neural progenitor identities. More broadly, the results suggest that primary regionalisation events coordinately govern multiple cell lineages at the cervicothoracic transition, with implications for understanding neurocristopathies involving combined enteric and trunk derivatives.

## Introduction

Neural crest cells (NCCs) represent one of the most critical progenitor cell types that underpins the development of the vertebrate body plan (Bronner & Simoes-Costa, 2016). Emerging transiently along the body axis, NCCs comprise four major subtypes that each arise at defined positions and give rise to characteristic derivatives: cranial, vagal, trunk, and sacral NCCs. In the cervicothoracic region, vagal NCCs make essential contributions to the developing heart (Kirby et al., 1983; Chan et al., 2004; Hutchins et al., 2019), in addition to neurons and glia within the enteric nervous system (ENS) (Le Douarin & Teillet, 1973; Hutchins et al., 2019). By contrast, trunk NCCs in this region produce sensory and sympathoadrenal cell types within the peripheral nervous system (Le Douarin & Teillet, 1975; Ma et al., 1996; Fraser and Bronner-Fraser, 1999). The crucial role of NCC subtypes is underscored by the diversity of congenital anomalies they affect. Such neurocristopathies (Bolande RP, 1997; Vega-Lopez et al., 2018) present regional impacts spanning both vagal and trunk derivatives, or more restricted effects (Croaker et al., 1998; Cheng et al., 2015). To resolve how such anomalies occur, it is essential to understand how NCCs acquire the right axial identity and how these mechanisms have evolved to coordinate the formation of the vertebrate body (Green et al., 2017).

At the head-to trunk transition, vagal NCCs emerge from multiple axial positions. While anterior vagal NCCs (somite levels 1 to 4 in mouse) can generate cardiac or enteric derivatives depending on their environment (Tang et al., 2021; Chan et al., 2004, Simkin et al., 2019; Espinosa-Medina et al., 2017), this plasticity is not shared with posterior vagal NCCs (somite levels 5 to 7 in mouse) which instead contribute to the ENS (Espinosa-Medina et al., 2017; Simkin et al., 2019), even following transplantation into anterior positions (Kuo and Erickson, 2011). Differences in cranial versus trunk NCC differentiation potential have been linked to distinct gene regulatory networks (GRNs) (Simoes-Costa & Bronner, 2016; Soldatov et al., 2019). Yet, whether such a model can fully account for developmental differences spanning the vagal region remains unclear (Soldatov et al., 2019; Kuo & Erickson, 2011, Simkin et al., 2019; Espinosa-Medina et al., 2017). Studies in avians (Ling & Sauka-Spengler, 2019; Krispin et al., 2010) and most recently in human (Vong et al., 2026) support the view that NCCs can undergo early fate restrictions, and derive from distinct lineages (Erickson et al., 2025; Lukoseviciute et al., 2023). Defining intrinsic differences, and how they are established across individual NCCs may shed light on the developmental mechanisms that result in congenital disease.

A potential mechanism that may contribute to regional differences spanning vagal NCCs is primary regionalization (Metzis et al., 2018). In this model, cells adopt discrete regional identities in the epiblast state (Simon & Hadjantonakis, 2018; Needham & Metzis, 2022). Primary regionalization depends on the activity of caudal type homeobox (CDX) transcription factors (TFs), which play a key role in controlling chromatin accessibility (Metzis et al., 2018). As CDX TFs are expressed in the caudal epiblast during late gastrulation, primary regionalization events precede the end of NCC specification along the body axis. CDX TFs are highly conserved across multiple animals, and their genetic removal results in NCC defects (Sanchez-Ferras et al., 2016; Rocha et al., 2021). Yet, to what extent primary regionalization impacts the variety of NCCs produced remains incomplete. As CDX TFs are partially redundant and their removal severely compromises embryo development (Young et al., 2009; van Rooijen et al., 2012), alternative strategies are needed to address this question.

In this study, we investigated how primary regionalisation of the epiblast contributes to the generation of NCCs in mouse embryos. Using in vivo mouse genetics and quantitative stem cell models of development, we show that a transient *Cdx1* lineage contributes to a subset of vagal neural crest cells and to trunk-to-tail derivatives, but that these related contributions are established through temporally distinct mechanisms. Single-cell spatial transcriptomics further resolves the molecular identities and overlapping distributions of individual vagal and trunk NCCs *in vivo*. Despite their close spatial relationship, vagal and trunk NCCs are specified by temporally and molecularly distinct programmes in which individual CDX paralogues establish different HOX states. Together, our findings suggest that distinct epiblast states preconfigure the identities of both neural crest and neural progenitors that subsequently emerge across the head-to-trunk transition.

## Results

### A *Cdx1* lineage contributes to posterior vagal NCCs

CDX TFs display a nested expression pattern in the embryo. *Cdx1* is detected around E7.25, followed by *Cdx2* and *Cdx4* (Meyer & Gruss, 1993; Gaunt et al., 2005). To test the extent to which CDX TFs contribute to the body plan, without disrupting embryogenesis, we engineered a tamoxifen-inducible *Cdx1^Cre-ERT2/+^* mouse line (Figure 1A; see methods) to perform lineage tracing. To test the specificity of the Cre relative to the expression of CDX1 expressing cells that could be tracked with the FLAG epitope, we crossed heterozygous mice to a well-established *R26R^tdTomato/+^* reporter mouse line (Madisen et al., 2010). Resulting *Cdx1^Cre-ERT2/+^; R26R^tdTomato/+^*(herein referred to as *iCdx1-Tdt*) mutant embryos harvested at E7.75, following exposure to tamoxifen at E6.25, prior to the onset of *Cdx1* (Meyer & Gruss, 1993) were subject to tissue immunofluorescence (Figure S1A-B). FLAG positive signal was detected in the caudal epiblast, consistent with the reported expression pattern of *Cdx1* (Meyer & Gruss, 1993), together with TdTomato fluorescence and nuclear ER staining, indicative of Cre activity (Figure S1A-B). Taken together, the generation of *iCdx1-Tdt* embryos enables lineage tracing of a *Cdx1* progenitor pool arising during late stages of gastrulation.

**Figure 1:**
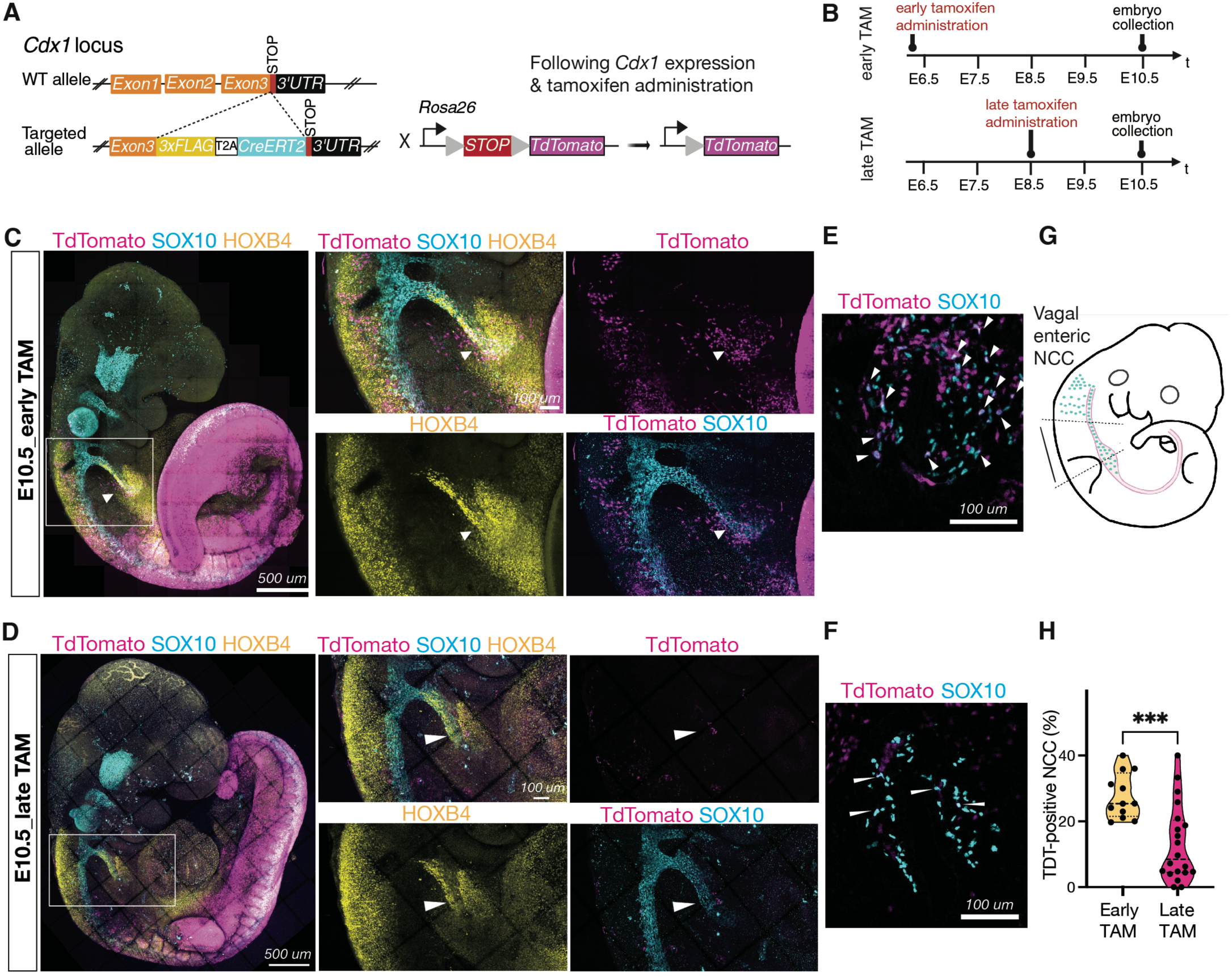
A *Cdx1* lineage transiently contributes to a subset of vagal NCCs. (A) Schematic of the *Cdx1* lineage tracing strategy to track Cdx1 positive epiblast progenitors using a Cdx1^Cre-ERT2/+^line inducible mouse line (see methods) crossed to the R26R^tdTomato/+^ reporter *(Madisen et al., 2010)*. (B) Schematic of the experimental design to test the contribution of early (Early TAM) versus late (Late TAM) Cdx1 expression. (C) WMIF of an E10.5 embryo (33 somites) shows *Cdx1* lineage contribution to vagal NCCs and trunk tissues. Magnified region (white box) shows the cervical region between somite 1 and 4. White arrows highlight co-localization of HOXB4 (yellow), TdTomato (magenta), and SOX10 (cyan). (D) WMIF of an E10.5 embryo (34 somites) shows the absence of TdTomato (magenta)/SOX10 (cyan) cells along the HOXB4-positive (yellow) vagus nerve (white arrow in the boxed magnified region). (E) Transverse section at the posterior vagal level (somite 5/6) shows co-localization of TdTomato (magenta) with NCCs contributing to dorsal root ganglia (DRG; labelled by SOX10 in cyan) and sympathoadrenal (SA) cells (expressing SOX10 and PHOX2B in yellow), sympathoenteric (SE, expressing SOX10/PHOX2B). White box indicates magnified view for enteric NCCs at the level of the foregut. (F) Transverse section at the forelimb level shows TdTomato (magenta) still contributes to DRG and SA (labelled by SOX10 in cyan) derivatives, while the contribution to enteric NCCS (white box and magnified view) is reduced (white arrows) upon delayed tamoxifen administration. (G-H) Quantification of the proportion of TdTomato-positive SOX10-NCC (H) in transverse section from the trachea to the future stomach region in embryos as indicated in G, exposed to tamoxifen (TAM) at the indicated times (B). WMIF, wholemount immunofluorescence; E, embryonic day; NCCs, neural crest cells; DRG, dorsal root ganglia; SA, sympathoadrenal; SE, sympathoenteric. All schematic were created with BioRender.com. Data are represented as mean ± SEM. *p value < 0.05, **p value < 0.01, ****p value < 0.0001.

To examine the contribution to NCCs along the rostrocaudal axis, *iCdx1-Tdt* embryos were collected at E10.5 and subject to wholemount immunofluorescence for SOX10 (Shibata et al., 2010). Tdt labelling was detected throughout the trunk and tail, in addition to partial labelling within cervical derivatives (Figure 1B-C). The *iCdx1-Tdt* lineage was detected in NCCs labelling the vagus nerve (white arrow, Espinosa-Medina et al., 2017), in contrast to no-tamoxifen controls (Figure S1C-D), suggesting that the *Cdx1* lineage contributes to vagal NCCs (Figure 1B-C), in addition to more caudal derivatives. Consistent with this view, *iCdx1-Tdt* embryos labelled vagal NCCs present between somite 3 and 7 at E9.25 (Figure S1E-G), with increasing contributions noted at somites 5-7, the axial position at which vagal NCCs give rise to enteric derivatives (Durbec et al., 1996; Chan et al., 2004; Espinosa-Medina et al., 2017).

We asked whether the contribution to vagal NCCs could be uncoupled from more caudal derivatives by controlling the timing of labelling. Delaying tamoxifen administration to E8.5 (Figure 1B; “late TAM”) no longer labelled cervical derivatives, including the vagus nerve, while labelling of trunk NCCs was maintained (Figure 1D). Quantifications of *iCdx1-Tdt* transverse sections confirmed that changing the timing of recombination significantly reduced the percentage of *iCdx1-Tdt* positive NCCs detected in mutliple transverse sections of the foregut (Figure 1E-H). Taken together, these results demonstrate that a *Cdx1* lineage arising during gastrulation transiently contributes to a subset of vagal NCCs that generate enteric derivatives and this contribution is temporally separable from its later role in generating trunk derivatives.

### Single-cell spatial transcriptomics resolves vagal and trunk NCCs *in vivo*

The partial contribution of a *Cdx1* lineage to vagal NCCs raises the possibility that vagal NCCs display discrete regional identities. Previous scRNA-seq analyses from NCCs isolated from mouse embryos also suggests that vagal and trunk NCCs are transcriptionally similar, despite yielding distinct derivatives (Soldatov et al., 2019; Liu et al., 2026). To resolve vagal versus trunk NCCs, we performed single-cell spatial transcriptomics at early migratory stages spanning a ∼36 hours window from E9.0 to E10.5 (Figure 2A). We captured axial positions encompassing both vagal enteric and trunk derivatives (somite 7-8) (Durbec et al., 1996; Espinosa-Medina et al., 2017); enteric (somite 5-6) (Durbec et al., 1996; Espinosa-Medina et al., 2017); cardiac and enteric (somite 1-4) (Chan et al., 2004; Durbec et al., 1996; Espinosa-Medina et al., 2017); or trunk NCCs (somite 9-10) (Durbec et al., 1996). Unsupervised clustering of all cells passing quality controls (see methods) recapitulated the expected cellular composition (Figure 2B, S2A-B), including neural progenitors, paraxial mesoderm and putative NCCs expressing *Sox10 (*Figure S2B) captured in ventral and dorsolateral migratory positions (Figure 2D).

**Figure 2:**
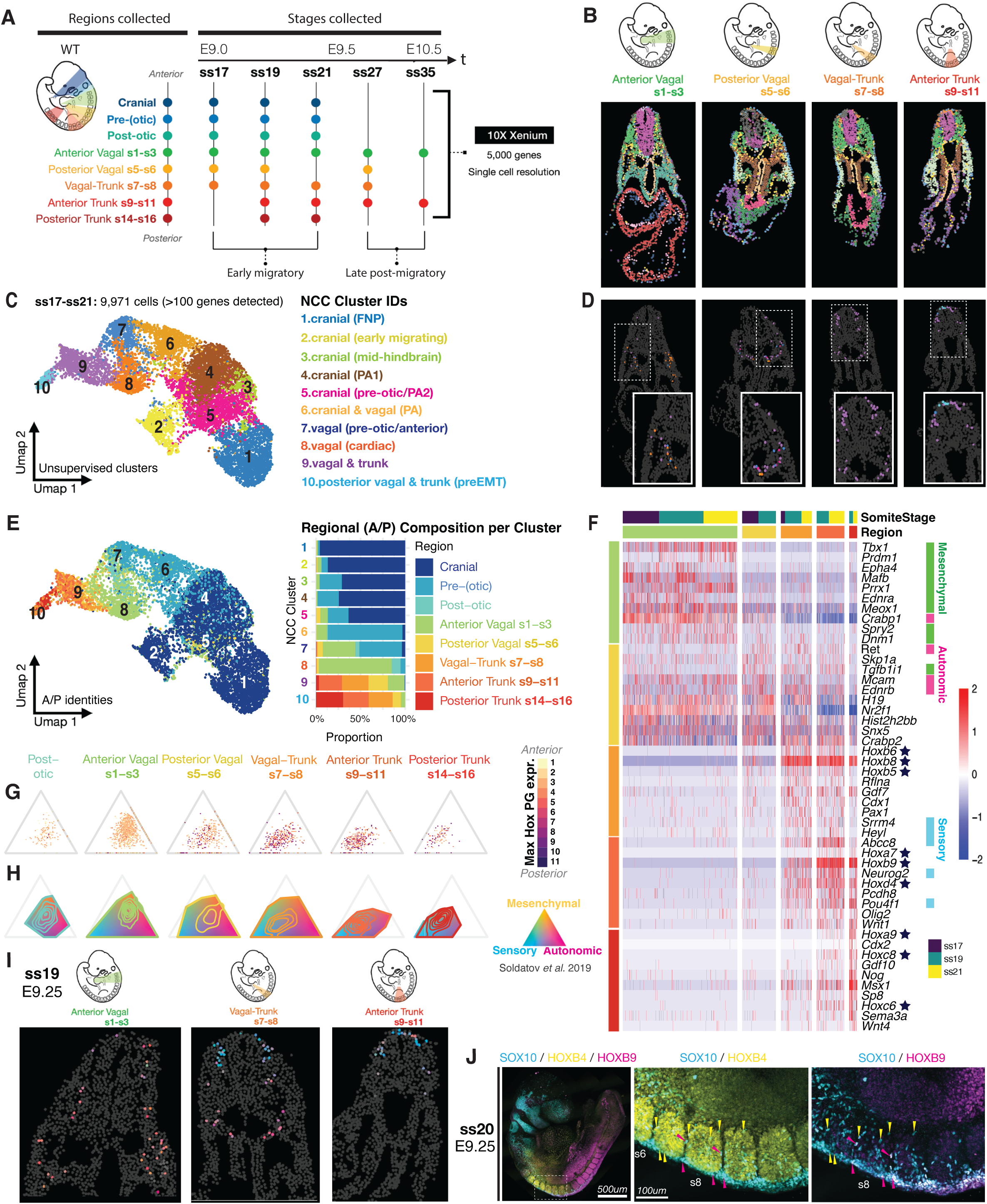
Single-cell spatial transcriptomics resolves a vagal-to-trunk neural crest transition *in vivo*. (A) Experimental design showing the stages and levels collected for spatial transcriptomics using the Xenium 10X platform with the pre-designed panel covering 5,000 gene targets. (B) Spatial annotation at the indicated level of a ss19 embryo showing the distinct clusters obtained after unsupervised clustering with NCCs in orange. (C-E) UMAP (C, E) and spatial (D) projections of annotated unsupervised clusters generated from NCC subsets extracted from ss17, ss19, and ss21 datasets (see Fig. S2A). Clusters in (E) are grouped by anatomical region, with regional composition shown in the bar plot. (F) Heatmap for the top 10 DEGs between anterior vagal (green), posterior vagal (yellow) and trunk (red) levels across the distinct time points ordered by log-fold change per level. (G-H) NCCs projected into a transcriptional landscape defined by sensory, autonomic, and mesenchymal competing gene modules (Soldatov et al., 2019). G shows the landscape coloured by maximum Hox paralogue group (PG1–PG11) identity. H shows the occupancy of state space for the indicated axial levels. Contours indicate cell density and outlines denote regional occupancy boundaries. (I) Spatial projection of the sensory–autonomic–mesenchymal transcriptional landscape shown in (G) in representative cervicothoracic NCC populations. Cells are coloured according to the relative contribution of sensory (cyan), autonomic (magenta), and mesenchymal (yellow) competing gene modules, with intermediate colours indicating mixed transcriptional states. (J) WMIF of an E9.5 (ss20) embryo shows co-existence of NCCs (stained with SOX10 in cyan) expressing either HOXB4 (in yellow, yellow arrows) or HOXB9 (in magenta, magenta arrows) in the region shown in the magnified box. ss, somite stage; UMAP, Uniform Manifold Approximation and Projection; DEGs, differentially expressed genes; NCC(s), neural crest cell(s); s, somite; E, embryonic day; WMIF, whole-mount immunofluorescence; IF, immunofluorescence.

To assess early NCC heterogeneity in the cervicothoracic region, we first verified that our limited number of targets (Xenium 5k panel) was capturing NCC diversity, by re-analysing published scRNA-seq data (Soldatov et al., 2019) with this reduced panel. 5k-derived clusters aligned well with original clusters (Figure S2C-D) and projection of NCC states on the 5k-derived UMAP revealed preservation of global manifold structure (Figure S2E), confirming that the 5k panel can be used to identify major NCC states. We, next integrated a total of 9,971 NCCs, each with at least 100 detected probes, extracted from the E9.0-E9.25 datasets (Figure S2A) and subjected them to unsupervised clustering. This revealed 10 clusters comprising distinct NCC states including early migratory progenitors (*Foxd3*/*Zic1*, Figure 2C and S2F) and differentiated NCCs with pharyngeal and mesenchymal identities (*Dlx2/Twist1*, Figure 2C and S2F). Projecting the anteroposterior identity of origin onto this UMAP showed the mixed regional composition of the early progenitor clusters (Figure 2E), showing the transcriptional similarities between posterior vagal and trunk NCCs as observed in other datasets (Figure S2G, Liu et al., 2026).

To resolve further NCC identity across vagal and trunk levels, we performed differential gene expression analysis. This revealed that posterior vagal and trunk NCCs adopt distinct autonomic (*Ret, Ednrb*) versus sensory (*Neurog2, Pou4f1*) signatures (Figure 2F) in line with their respective sympathoenteric, dorsal root and sympathetic ganglia contribution (Espinosa-Medina et al., 2017; Durbec et al., 1996; Ma et al., 1999). Along these autonomic and sensory signatures, distinct *Hox* genes expressed in vagal versus trunk levels were detected: posterior vagal NCCs are enriched with a cervical *Hox* identity (*Hoxb4-8*; Howard et al., 2021, 2022) while trunk NCCs express *Hoxb9* (Figure 2F) (Parker et al., 2016; Mallo et al., 2010). Similar differences in *Hox* genes separating vagal from trunk NCCs at E9.0 were also detected at E10.5 in derivatives arising from such NCCs. In particular, sympathoenteric derivatives displayed *Hoxb4-8* expression while DRG/SG expressed *Hoxb4-9* (Figure S2H-I). Taken together, these data show that differences in thoracic *Hox* expression delineate vagal versus trunk NCCs.

After defining differences in NCC identity, we next sought to understand how they were structured in the cervicothoracic region. To test how the co-activation of previously observed competing programs for sensory, autonomic and mesenchymal fates (defined by Soldatov et al., 2019) was spatially distributed, we assessed their co-expression at each axial position. Co-detection of autonomic and mesenchymal modules was observed at anterior vagal positions (Figure 2G) (Soldatov et al., 2019), while posterior vagal and trunk levels displayed co-activation of sensory and autonomic modules which coincides with an upregulation in *Hoxb9* expression (Figure 2G). Strikingly, at the somite 7-8 level, neighbouring cells with sensory versus autonomic modules were observed (Figure 2G-H). Immunofluorescence to examine HOXB4 and HOXB9 signal intensities in NCCs (Figure 2I) further verified that at the vagal-to-trunk transition (somite 7-8), NCCs expressed either HOXB4 or HOXB9, indicating distinct NCC subtypes detected at the single cell level within the same axial position. Taken together, somite 7-8 defines a transitional zone harbouring both posterior vagal and trunk NCCs.

### CDX transcription factors are required for NCC regionalization

Since vagal and trunk NCCs are captured in the same axial location *in vivo*, how do NCCs adopt vagal or trunk states? CDX TFs are well-established regulators of the cervicothoracic transition, that play partially redundant roles in controlling *Hox* genes (Van den Akker et al., 2002; Young et al, 2009; Subramanian et al., 1995; Amin et al., 2016). To address how CDX TFs impact vagal and trunk identities, we developed a quantitative *in vitro* model of NCCs from mouse ESCs, benchmarked against the corresponding NCC states resolved from our single-cell spatial transcriptomics data from mouse embryos. Efficient generation of cranial NCCs *in vitro* from human (h)ESCs relies on a precise balance of active WNT and BMP signaling (herein referred to as WNT/BMP), applied to cells that have reached an epiblast-like state (Hackland et al., 2017; Frith et al., 2018; Figure 3A). Applying these principles to generate mouse cranial NCCs *in vitro*, we differentiated mESCs over a three-day period to generate anterior epiblast-like (EpiL) progenitors using established methods (Gouti et al., 2014; Metzis et al., 2018; Amblard et al., 2025), followed by NCC induction with WNT/BMP (Figure S3A-D) (Hackland et al., 2017; Frith et al., 2018). Within 24 hours of exposure to NCC inducing cues, NCC-like progenitors expressing TFAP2A (Figure S3B-C) were detected (day 4; 71.7%), followed by SOX10, 48 hours later (day 5; 82.6%) (Figure 3A; S3B-C) recapitulating observations from hESCs (Hackland et al., 2017; Frith et al., 2018).

**Figure 3:**
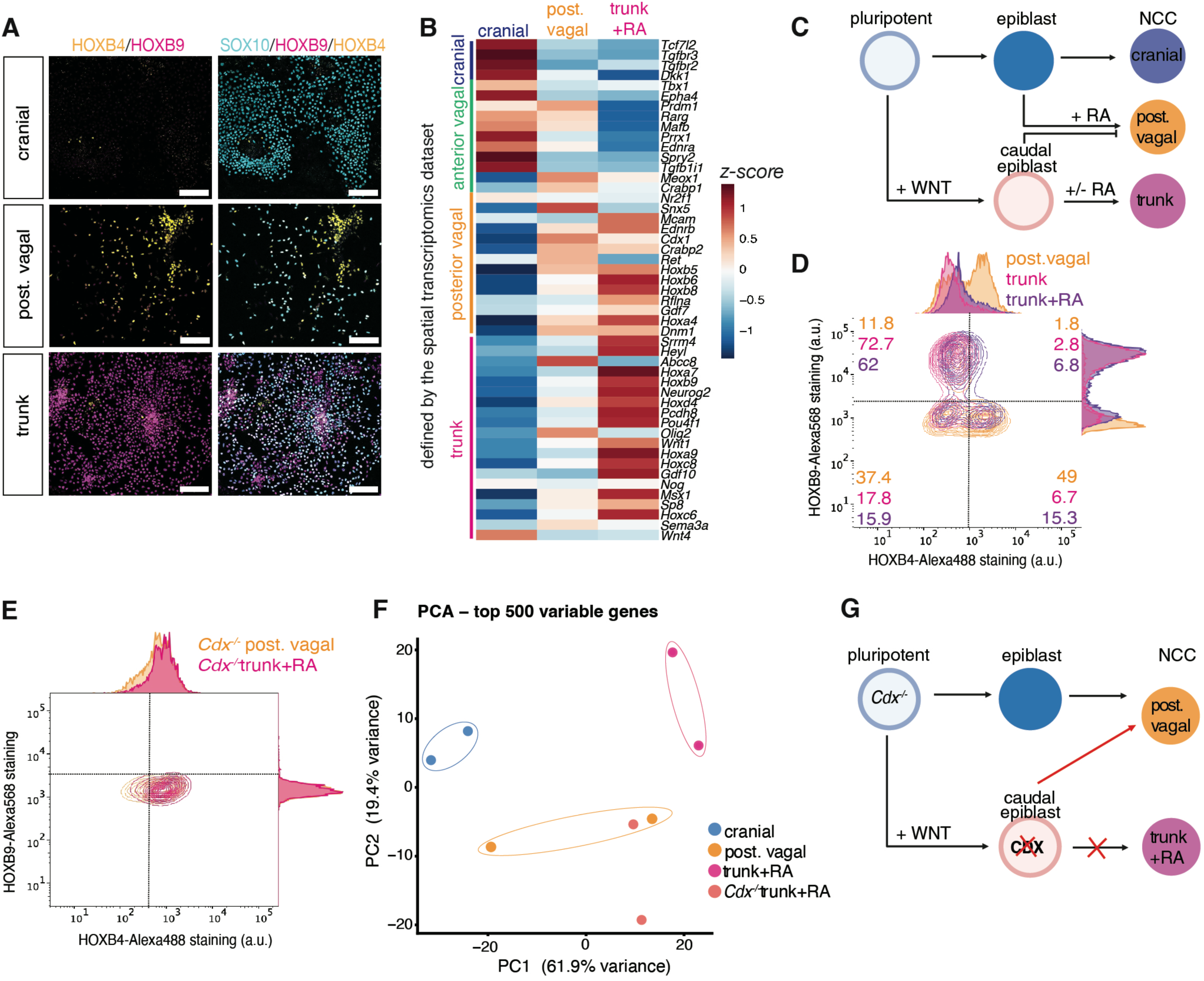
Removal of CDX factors derepresses a vagal NCC programme *in vitro*. (A) IF demonstrates SOX10-positive NCC-like progenitors (cyan) expressing HOXB4 (yellow) in the posterior vagal condition versus HOXB9 (magenta) in the trunk NCC condition. (B) Heatmap showing the expression of genes identified *in vivo* that define anterior vagal, posterior vagal or trunk NCCs. (C) Schematic summarizing the NCC subtypes produced *in vitro* from mouse ESCs. (D-E) Representative flow cytometry plots for HOXB4/9 showing TFAP2A-gated NCCs generated from WT (D) cells express either HOXB4 (posterior vagal in orange) or HOXB9 (trunk and trunk+RA in magenta and purple) while NCCs generated from Cdx^-/-^ (E) cells (Gouti et al., 2014) express only HOXB4 (in posterior vagal and trunk+RA conditions). (F) Dimensionality reduction using principal component analysis shows Trunk+RA NCCs generated from Cdx^-/-^ cells resemble more *WT* Posterior Vagal than *WT* Trunk+RA NCCs. (G) Schematic summarizing findings from Cdx^-/-^ mESCs. All schematics were created with BioRender.com. *p value < 0.05, **p value < 0.01, ****p value < 0.0001. NCC(s), neural crest cell(s); mESCs, mouse embryonic stem cells; ATRA, all-trans-retinoic-acid; IF, immunofluorescence.

We next tested the capacity to modulate the regional identity of NCCs produced *in vitro*. Retinoic acid (RA) signaling regulates the cervicothoracic Hox code (Bel-Vialar et al., 2002; Ishikawa and Ito, 2009) and promotes the generation of vagal NCCs when applied to cranial NCCs generated from hESC (Fattahi et al., 2016; Frith et al., 2018, 2020). Similarly, addition of all-trans RA (ATRA) between day 4-5 resulted in the generation of posterior vagal NCC-like progenitors expressing HOXB4 (Figure 3A). Consistent with previous reports (Frith et al., 2020), thoracic *Hox* genes such as *Hoxb9* were not induced by ATRA exposure (Figure 3A-B), suggesting the generation of vagal NCCs was not accompanied by trunk NCCs. As observed in hESC (Frith et al., 2018; Fan et al., 2023), efficient generation of trunk NCCs can be produced from a caudal-epiblast-like (CEpiL) progenitor state generated by exposing epiblast progenitors to bFGF/CHIR to promote active WNT signalling conditions (Figure S3D). Similarly, we found that the CEpiL progenitors exposed to BMP/WNT no longer produced HOXB4 and instead, generated HOXB9-positive NCCs (Figure 3A, Frith et al., 2018; Fan et al., 2023). To assess the regional identity of NCCs produced in cranial, vagal or trunk conditions, mRNA-seq was performed at the day 5 timepoint, confirming the expression of a NCC programme (Figure S3F). Region-specific genes identified in the spatial transcriptomics dataset generated from mouse embryos revealed discrete signatures corresponding to these states, as demonstrated by the expression of *Tgfbr2/3, Dkk1* (cranial, Soldatov et al., 2019; Conway & Kaartinen, 2011), *Sox9, Ret* (posterior vagal) or *Neurog2, Pou4f1, Msx1, Sp8* (trunk) (Figure 3B). These conditions also recapitulated the *Hox* states that define posterior vagal (*Hoxb5/6/8*, *Hoxa4*) versus trunk NCCs (*Hoxb9, Hoxa7, Hoxa9, Hoxc8*) (Figure 3B), providing us with a system to test how these defined regional *Hox* states are established.

Since CEpiL progenitors transiently express *Cdx1/2/4* (Figure S3D-E), we next asked whether CEpiL cells can generate vagal as well as trunk NCCs, depending on exposure to extrinsic cues such as RA. To test this, we exposed CEpiL progenitors to NCC inducing cues including ATRA (referred to herein as trunk+RA). (Figure 3C-D). Strikingly, CEpiL cells exposed to RA are not capable of efficiently generating vagal NCCs and instead maintain a trunk *Hox* code, with 68.8% of TFAP2A-gated NCCs expressing HOXB9, while a reduced proportion of cells expressed HOXB4 (22.1%). Taken together, NCCs derived from a CEpiL state adopt a trunk identity, independent of their exposure to RA signaling cues.

To determine how exposure to WNT signaling enables the generation of trunk NCCs, we tested the role of CDX TFs, which are essential in controlling the CEpiL state and have been shown to play a role in the generation of trunk NCCs (Frith et al., 2018; Fan et al., 2023; Lukoseviciute et al., 2023). As CDX ablation compromises trunk development (Amin et al., 2016), we took advantage of our *in vitro* assay to probe the regional identity of NCCs produced from *Cdx*-lacking mESCs (*Cdx ^-/-^*, Gouti et al., 2014). Under trunk+RA NCC conditions, (Figure 3E), *Cdx* mutant cells still generate TFAP2A-expressing NCCs but no longer adopted a trunk identity as previously shown (Gouti et al., 2014; Gogolou et al., 2022), and instead expressed HOXB4 (Figure 3E). To probe the variance present across control versus *Cdx*-lacking NCCs, we generated mRNA-seq data from control versus *Cdx ^-/-^* NCCs in trunk conditions with RA present. Principal component analysis (see methods) revealed that trunk+RA *Cdx^-/-^* NCCs grouped more closely with posterior vagal NCCs (Figure 3F). Thus, CDX TFs repress the generation of vagal NCCs *in vitro* (Figure 3G).

### Distinct CDX paralogues control NCC subtype identity

Despite sharing a lineage, combined activity of CDX transcription factors represses a vagal NCC state. To reconcile how a *Cdx1* lineage transiently generates vagal NCCs, we tested the hypothesis that individual CDX factors may perform separate roles. CDX1 and CDX2 play partially redundant roles *in vivo* (Calon et al., 2007; Grainger et al., 2013; Domon-Dell & Freund, 2002; Savory et al., 2009). *Cdx1* also precedes *Cdx2/4* expression *in vivo* (Meyer and Gruss, 1993; Beck et al., 1995), an effect also recapitulated *in vitro* (Mariani et al., 2021; Gouti et al., 2014; Blassberg et al., 2022). We hypothesized that the timing of expression of *Cdx1* versus *Cdx2* may ensure the generation of vagal derivatives in a limited and temporal manner (Figure 1B-C). To test this model, we reduced the duration of active WNT signaling conditions to a 12h-pulse to attempt to capture the onset of CDX1 prior to CDX2 (Figure 4A). This short exposure resulted in CDX1-positive EpiLCs lacking CDX2 (referred to as early-CEpiLCs, Figure 4B). Early-CEpiLCs subjected to NCC inducing cues generated vagal NCCs expressing HOXB4 (Figure 4C) and lacking HOXB9, unlike the longer exposure provided to CEpiLCs which promotes a shift to a trunk HOXB9 identity (Figure 4D). This suggests that altering the duration of WNT activating conditions in EpiLCs uncouples the production of vagal from trunk derivatives (Figure 1B-C).

**Figure 4:**
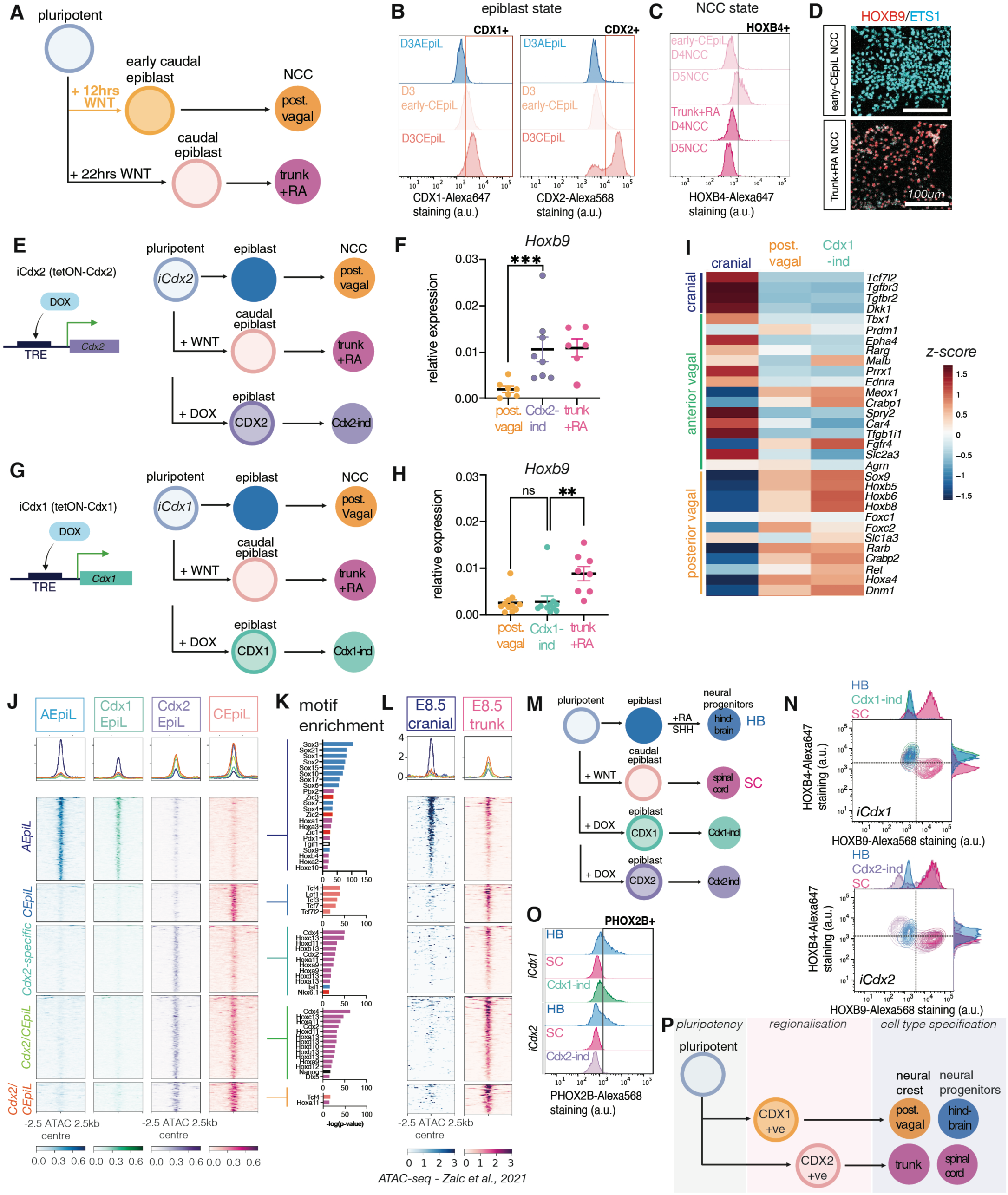
Separate epiblast regionalisation events dictate neural crest and neural progenitor identity. (A) Schematic summarising the conditions and time points assayed (B-D) to test the impact of a shortened pulse (12 hours) of active WNT signalling conditions provided in the epiblast state. (B) Cytometry histogram for CDX1 and CDX2 showing early-CEpiL (light pink) express CDX1 while CEpiL (pink) express both CDX1 and CDX2. (C) Cytometry histogram for HOXB4 of NCCs generated from Early-CEpiL or CEpiL shows only TFAP2A-gated Early-CEpiL NCCs express HOXB4. (D) IF analysis of NCC generated from Early-CEpiL or CEpiL shows ETS1 (cyan) cells derived from Early-CEpiL do not acquire a trunk (HOXB9; red) identity. (E-H) Schematic of the Tet-ON inducible cell lines generated to control the induction of *Cdx2* (*iCdx2, E*) and *Cdx1* (*iCdx1*, G), simplified schematic of the 5-day NCC differentiation conditions (E,G) and relative expression (RT-qPCR) for *Hoxb9* in NCC progenitors generated from *iCdx2* cells (Cdx2-ind) shows the induction of *Cdx2* is sufficient to upregulate *Hoxb9* (F), unlike *Cdx1* induction (H, Cdx1-ind). (I) Heatmap showing *Cdx1* induction (Cdx1-ind) promotes a posterior vagal NCC identity indicated by expression of target genes detected *in vivo*. (J) Heatmap showing ATAC-seq coverage for elements differentially accessible between any two conditions with the same dynamics grouped in clusters, shows opening of shared Cdx2/CEpiL or Cdx2- and CEpiL-specific sites while AEpiL-specific sites are closed in an incremental manner in Cdx1EpiL, Cdx2EpiL and CEpiL. (K) Motif enrichment analysis for the elements reveals motifs for posterior HOX and CDX in the shared Cdx2/CEpiL clusters. (L) Heatmap showing ATAC-seq coverage for the same elements in cranial (blue) or trunk (pink) OCT4-positive NCCs isolated from E8.5 embryos (Zalc et al., 2021). (M) Simplified schematic of the 5-day neural progenitor (NP) differentiation from mESCs. (N) Representative flow cytometry plots for HOXB4/9 showing SOX1-gated NP generated from Cdx1EpiL (Cdx1-ind, green) express HOXB4 while SOX1-gated NP generated from Cdx2EpiL (Cdx2-ind, purple) express HOXB9. (O) Cytometry histogram for PHOX2B of NP shows Cdx2 induction in EpiL (purple) conditions represses PHOX2B unlike *Cdx1* induction (green). (P) Model for cervicothoracic diversification: epiblast cells express Cdx1 and can give rise to posterior vagal NCCs or caudal hindbrain progenitors. As Cdx1 progenitors progress towards a Cdx2 positive state, a trunk identity is promoted in NCCs and neural progenitors. All schematics were created with BioRender.com. Data are represented as mean ± SEM. *p value < 0.05, **p value < 0.01, ****p value < 0.0001. NCC(s), neural crest cell(s); mESCs, mouse embryonic stem cells; CEpiL, caudal epiblast-like; EpiL, epiblast-like; AEpiL, anterior epiblast-like; NP, neural progenitors; E, embryonic day; D, day; SC, spinal cord; Hb, hindbrain.

To assess how the duration of active WNT signalling conditions in EpiLCs alters their capacity to produce vagal and trunk NCCs, we engineered doxycycline-inducible mESCs to test the role of *Cdx1* versus *Cdx2* (*iCdx1* vs *iCdx2*) in EpiLCs, independently of active WNT signalling conditions (Figure 4E,G). After doxycycline (DOX) addition between day 2-3, EpiLCs were detected expressing CDX1 (Cdx1EpiL) or CDX2 (Cdx2EpiL) at comparable protein levels to the level detected in CEpiL progenitors that express both paralogues, in addition to other WNT-induced target genes (Figure S4A-D). Strikingly, the induction of *Cdx2* generates HOXB9-positive trunk NCCs (Cdx2-ind, Figure 4F; S4E) in a similar manner to CEpiL conditions. By contrast, the induction of *Cdx1* alone produced NCCs lacking *Hoxb9* and resembling posterior vagal NCCs (Cdx1-ind, Figure 4G-I). These data suggest that *Cdx1* and *Cdx2* independently impact the generation of posterior vagal (repressed by CDX2, Figure S4E) versus trunk (CDX2-driven) NCCs.

### *Cdx1* and *Cdx2* induce distinct chromatin landscapes

Having shown that *Cdx1* versus *Cdx2* induction uncouples the generation of vagal versus trunk NCCs *in vitro*, we next investigated the potential underlying molecular mechanism. Since CDX2 induction can promote chromatin accessibility changes (Metzis et al., 2018), we hypothesized that Cdx1EpiL and Cdx2EpiL cells display distinct chromatin landscapes. To test this, we performed ATAC-seq on AEpiL, Cdx1EpiL, Cdx2EpiL and CEpiL cells (Figure 4E,G). K-means clustering of differentially accessible sites detected across all four conditions revealed discrete chromatin landscapes corresponding to a set of AEpiL-specific regions, which loses accessibility in CEpiL progenitors; a set of Cdx-independent CEpiL-specific regions, and two shared *Cdx2*/CEpiL clusters (Figure 4J, S4G). *Cdx1* induction did not trigger an increase in chromatin accessibility at sites observed with Cdx2. Instead, decreased accessibility at AEpiL sites was observed upon *Cdx1* induction, with further reduction observed following CDX2 or CEpiL conditions (Figure 4J). Motif enrichment analysis confirmed that the AEpiL-associated sites were enriched with SOX/ZIC motifs, while the CEpiL-specific cluster was exclusively enriched in TCF/LEF motifs. CDX2/4 and HOX motifs defined the *Cdx2* or *Cdx2*/CEpiL shared sites (Figure 4K). To test whether these regional accessibility patterns were detectable in NCCs *in vivo*, we examined the accessibility patterns present in cranial versus trunk NCCs from mouse embryos (Zalc et al., 2021). Strikingly, cranial NCCs displayed accessibility at AEpiL sites. By contrast, CEpiL or *Cdx2*-dependent sites were accessible in trunk, but not cranial, NCCs (Figure 4L), in agreement with previous findings (Zalc et al., 2021). Taken together, differential chromatin landscapes established by *Cdx1* versus *Cdx2 in vitro* preconfigure epiblast progenitors with regionally distinct accessibility landscapes, enabling the generation of discrete subtypes of NCCs.

Since CDX1 and 2 promote distinct chromatin accessibility landscapes at the epiblast stage, their activity could influence the generation of additional cell types. In particular, neural progenitors, which adopt a similar Hox state to NCCs (Figure S4H-I) require primary regionalization events in the epiblast to produce thoracic spinal cord (Metzis et al., 2018). To test if CDX paralogue state acquired in the epiblast state impacts the generation of spinal cord, we used our *iCdx1* versus *iCdx2* ESCs to respectively generate CDX1 or CDX2 positive EpiL progenitors, followed by exposure to motor neuron progenitor-inducing cues, as previously performed (Gouti et al., 2014; Metzis et al., 2018) (Figure 4M). Following CDX2 activity, resulting neural progenitors adopted a HOXB9 identity, similar to spinal cord progenitors (Figure 4N-O). By contrast, the activity of CDX1 did not promote HOXB9, and instead, produced visceral motor neural progenitors expressing HOXB4 and PHOX2B (Figure 4N-O). In summary, these findings demonstrate that distinct epiblast states not only alter the subsequent identity of NCCs, but also apply to the generation of neural progenitors (Lippmann et al., 2015; Mouilleau et al., 2021) that produce distinct cell types spanning the head-to-trunk transition (Figure 4P).

## Discussion

Here, we tested the mechanisms that contribute to NCC subtype identity. Using mouse genetic lineage tracing, a single-cell resolved spatial transcriptomic atlas from mouse embryos, and *in vitro* models of mouse NCC and neural progenitor development, we identify a lineage that transiently produces posterior vagal NCCs together with trunk-to-tail derivatives. We provide evidence that, despite transiently sharing a common progenitor population, vagal and trunk NCCs are established through distinct epiblast regionalisation events governed by the timing and expression of CDX paralogues. These findings revise models of NCC and neural progenitor regionalisation at the cervicothoracic transition by suggesting that axial subtype identity is preconfigured before the emergence of NCC states.

Using mouse lineage tracing approaches, we identify a *Cdx1* progenitor population that contributes to a subset of vagal NCCs in a transient manner that is separable from trunk-to-tail derivatives. These findings raise the possibility that the transient developmental window in which posterior vagal and trunk NCCs share a lineage represents a point of vulnerability in neurocristopathies involving both enteric and trunk derivatives (Croaker et al., 1998; Gaultier et al., 2005; Nagashimada et al., 2012; Cheng et al., 2015). As the removal of *Cdx* genes disrupts enteric and trunk NCC derivatives (Sanchez-Ferras et al., 2016; Rocha et al., 2021), the timing of perturbation may contribute to the spectrum of phenotypes observed across different neurocristopathies (Vega-Lopez et al., 2018).

That a *Cdx1* lineage contributes in a partial, rather than complete, manner to the generation of vagal NCCs supports the view that vagal NCCs can arise from separate developmental origins (Durbec et al., 1996; Espinosa-Medina et al., 2017; Erickson et al., 2025). Consistent with this hypothesis, our spatial transcriptomics data further resolve distinct vagal NCC states *in situ*, supporting the idea that vagal NCCs represent a transitional population spanning the cervicothoracic region (Kuo & Erickson, 2011; Simkin et al., 2019; Tang et al., 2021). That individual NCCs display varying degrees of sensory, autonomic or mesenchymal states at a given position (Soldatov et al., 2019) also suggests that neighbouring cells exhibit transcriptional biases that may not be fully explained by their surrounding environment (Hsu et al., 2026). These spatial transcriptomic datasets provide a molecular framework for understanding neural crest heterogeneity, and the broader cellular diversity present at the head-to-trunk transition.

The close spatial and transcriptional similarity between posterior vagal and trunk NCCs prompted us to investigate how these distinct identities are controlled. We confirmed that, despite sharing a developmental lineage, NCCs spanning the cervicothoracic region adopt distinct molecular states dominated by *Hox* gene expression differences (Mallo et al., 2010; Parker et al., 2016; Duboule et al., 1989). Re-engineering NCC-like progenitors demonstrated that control over CDX expression in the epiblast state determines posterior vagal versus trunk identity *in vitro*. Individual and combinatorial CDX transcription factor expression established distinct chromatin accessibility landscapes, and these differences corresponded to regionally restricted programmes present in NCCs *in vivo* (Zalc et al., 2021).

Our findings suggest that CDX transcription factors establish distinct regional programmes in the epiblast state, with consequences that extend beyond NCC identity. We show that the regionalisation events that distinguish vagal and trunk neural crest subtypes also extends to the generation of neural progenitors. Thus, early regionalisation of shared progenitor populations may provide a general strategy for coordinating the production of multiple cell types at the head-to-trunk transition. More broadly, these findings support a model in which developmental diversity is shaped by epiblast regionalisation programmes that preconfigure the identity of descendant cell types. Defining how such regionalisation strategies operate across vertebrates will be important for understanding how these events coordinate cellular diversity.

## Author contributions

Conceptualisation: Irène Amblard (I.A.), Christos M Kalaitzis (C.M.K.), Juan M Vaquerizas (J.M.V.) and Vicki Metzis (V.M.). Data Curation: I.A., C.M.K. and Hothri Ananyambica Moka (H.A.M.). Formal Analysis: I.A., C.M.K. and H.A.M. Funding Acquisition: J.M.V. and V.M. Investigation: I.A., C.M.K., Ivan Andrew (Iv.A.) and Ka Lok Choi (K.L.C.). Methodology: I.A., C.M.K., Irina Balaguer Balsells (I.B.B.), Laurence Game (L.G.) and V.M. Project Administration: I.A. and V.M. Resources: J.M.V. and V.M. Software: C.M.K. and H.A.M. Supervision: I.A., L.G., J.M.V., V.M. Validation: I.A. Visualisation: C.M.K. and J.M.V. Writing – original draft: I.A., C.M.K. and V.M. Writing – review and editing: I.A., C.M.K., I.B.B., Iv.A., K.L.C., H.A.M., L.G., J.M.V. and V.M.

## Acknowledgements

We thank James Briscoe, Tom Frith, Tristan Rodriguez, and all lab members for comments on the manuscript. We are grateful to Robert Blassberg and James Briscoe for their support in generating the Cdx1 and Cdx2 Tet-ON mouse embryonic stem cells and targeting plasmids. For support, training and access to equipment, we thank James Elliott and Bhavik Patel from the LMS/NIHR Imperial Biomedical Research Centre Flow Cytometry Facility, the staff at the Central Biomedical Services unit at Imperial College London, as well as Chad Whilding and Dirk Dormann at the LMS microscopy facility. This work was supported by core funding to the Laboratory of Medical Sciences from the Medical Research Council and a Sir Henry Dale Fellowship awarded to V.M., jointly funded by the Wellcome Trust and the Royal Society (grant number 218536/Z/19/Z).

## METHODS

### Mouse work

All animal procedures were performed in accordance with the Animal (Scientific Procedures) Act 1986 under the UK Home Office project licenses PP6959042 and PBBEBDCDA and approved by the Imperial College London Animal Welfare and Ethics Review Body committee.

### _Cdx1_Cre-ERT2/+_; R26R_tdTomato/+ _mouse line_

The *Cdx1^Cre-ERT2/+^* mouse line was generated by Ozgene Pty Ltd (Australia). Gene targeting was performed in C57BL/6 ESCs to introduce a 3xFLAG tag followed by a T2A and Cre-ERT2 cassette immediately upstream of the endogenous STOP codon in the third exon of *Cdx1* (Fig 1A). Correctly targeted ESC clones were validated by PCR and Sanger sequencing. Heterozygous mice were viable, fertile and maintained on a C57BL/6J background. Genotyping was performed by PCR using the Hotshot protocol with primers listed in Table S1f. The Cre targeted allele amplified a product of 104 bps bp.

To perform lineage tracing, *Cdx1^Cre-ERT2/+^* mice were crossed with the *Gt(ROSA)26Sor^tm14(CAG-tdTomato)Hze^*(Jax stock number 007914; referred to as R26R^tdTomato/+^) to generate *Cdx1^Cre-ERT2/+^; R26R^tdTomato/+^* mice. Genotyping PCRs were performed as described above to genotype Cre heterozygous carriers. The *R26R^tdTomato/+^* allele was detected by PCR using primers listed in Table S1f and produced a product of 297 bps bp for the wildtype allele and 200 bps bp for the targeted allele.

*Cdx1^Cre-ERT2/+^; R26R^tdTomato/+^* mice were time-mated overnight. Vaginal plugs were checked every morning, and, if positive, the embryonic day was defined at E0.5. This was followed by oral gavage with 4-hydroxytamoxifen (tamoxifen, see Table S1a) at 0.1 mg/body weight dissolved in corn oil at indicated times. Embryos were collected at indicated times, dissected in EmbryoMAX M2 medium for E7.5 and ice-cold PBS for E9.5/E10.5, screened, and genotyped using the yolk sacs.

### Cell lines

All mouse ESC lines were maintained at 37 °C with 5% CO2 and routinely tested for mycoplasma contamination. All ESC lines used were derived from the XY HM1 line (Doetschman et al., 1987) or the Tet-ON (Serafimidis et al., 2008). The *iCdx1/2* cell lines were derived from the XY HM1 Tet-ON line (Serafimidis et al., 2008) by electroporating the HPRT locus targeting vector Hprt2 containing the *Cdx1* or *Cdx2* cDNA built using Sal1/Not1 restriction digest as previously described (Serafimidis et al., 2008).

Integrants were selected by culturing for 10 days in hypoxanthine-aminopterin-thymidine (HAT)-containing ESC medium. Following adaptation in 2i/LIF, clones were picked and expanded. Construct integration was confirmed by genotyping (see primers in Table S1e) and sequencing, and transgene expression was confirmed by imaging and RT-qPCR.

### ESC culture and differentiation

All mouse ESCs were expanded on mitotically inactivated mouse embryonic fibroblasts (feeders) in ESC medium (DMEM knockout medium supplemented with 1.000U/ml LIF, 10% cell-culture-validated fetal bovine serum, and 2mM L-Glutamine).

To differentiate mESCs into neural crest cells, mouse ESCs were differentiated in methods comparable to human ESCs using the Top-Down inhibition approach (Frith et al., 2018, 2020; Hackland et al., 2017). ESCs were dissociated with 0.05% trypsin, and plated on tissue-culture-treated plates for two sequential 15-20 minutes (mins) periods in ESC medium to separate them from their feeder layer cells, which adhere to the plastic. To start the differentiation, cells remaining in the supernatant were pelleted by centrifugation, counted, and resuspended in N2B27 medium containing 10 ng/ml bFGF, and 5x10^4^ cells per 35 mm gelatin-coated CellBIND dish or 6-well plate (Corning) were plated. N2B27 medium contained a 1:1 ratio of DMEM/F12:Neurobasal medium (Gibco) supplemented with 0.5% N2 (Gibco), 1% B27 (Gibco), 2mM L-glutamine (Gibco), 40mg/ml BSA (Sigma), and 0.1mM 2-mercaptoethanol.

To generate epiblast-like (EpiL) cells, the cells were grown respectively for 2 days in N2B27

+10 ng/ml bFGF. To generate anterior epiblast-like (AEpiL), EpiL were grown respectively for an extra day in N2B27 +10 ng/ml bFGF + dual SMAD-inhibitors (10uM SB + 2uM DMH1). To generate caudal epiblast-like (CEpiL) cells, EpiL were cultured in N2B27 + 10 ng/ml bFGF +5 μM CHIR99021 + dual SMAD-inhibitors for a day (22hrs).

To generate early caudal epiblast (early-CEpiL) cells, EpiL were cultured in N2B27 + 10 ng/ml bFGF +5 μM CHIR99021 + dual SMAD-inhibitors for 12hrs before replacing the media with NCC media (DMEM/F-12 medium containing 0.5% N2, 40mg/ml BSA (Sigma), and 0.1mM 2-mercaptoethanol, 1% GlutaMAX) supplemented with 15ng/ml recombinant BMP4, 1uM CHIR99021, and dual SMAD-inhibitors (2uM SB + 1uM DMH1) for an extra 10hrs.

To generate single Cdx1-or Cdx2-EpiL cells, EpiL cells were cultured in N2B27 + 10 ng/ml bFGF + 1ug/uL Doxycycline + dual SMAD-inhibitors for a day.

To generate neural crest cells with distinct axial identities, AEpiL, CEpiL, early-CEpiL or Cdx1-and Cdx2-EpiL progenitors were cultured in NCC media (DMEM/F-12 medium containing 0.5% N2, 40mg/ml BSA (Sigma), and 0.1mM 2-mercaptoethanol, 1% GlutaMAX) supplemented with 15ng/ml recombinant BMP4, 1uM CHIR99021, and dual SMAD-inhibitors (2uM SB + 1uM DMH1) for two days. Media was refreshed every day from day 3-5 for all experiments.

To generate vagal neural crest cells, progenitors with distinct identities (AEpiL, CEpiL, early-CEpiL or Cdx1- and Cdx2-EpiL) were cultured in NCC media supplemented with 15 ng/uL BMP4, 1uM CHIR99021, and dual SMAD-inhibitor (2uM SB + 1uM DMH1). After one day of culture, media were replaced with the same NCC media supplemented with 15 ng/uL BMP4, 1uM CHIR99021, dual SMAD-inhibitor (2uM SB + 1uM DMH1), and ATRA 100nM for one day.

### Wholemount immunofluorescence

Mouse embryos from time-mated pregnant females were collected at E10.5 and fixed in 4% PFA for 2hrs at 4°C under gentle agitation. After PBS washes, embryos were permeabilized for 45mins in PBS + 0.5% Triton-X100 before incubation with PBS + 0.1% Triton-X100 + 2% BSA + 4% donkey serum for blocking for at least 2hrs at room temperature (RT) under gentle agitation. Embryos were then incubated overnight at 4°C under gentle agitation with primary antibodies in blocking media before extensive washes in PBST (PBS + 0.1% Triton-X100) at RT under gentle agitation for a whole day. Embryos were incubated overnight at 4°C in blocking media before incubation with secondary antibodies (see Table S1i) in the dark at RT. After further washes with PBST, embryos were incubated for 1hr in PBST with Dapi (1/10000) before washing with PBST and PBS at RT. For imaging, embryos were mounted in 1.5% low-melt agarose into glass-bottom Ibidi 35mm dishes. Images were acquired on a Leica SP8 DLS. Used primary antibodies are listed in Table S1b.

### Tissue sectioning and immunofluorescence

Mouse embryos from timed pregnant females were collected and fixed in 4% PFA for 1.5 to 2hrs at 4°C under gentle agitation, washed in PBS, and transferred to 30% sucrose in PB (0.12M phosphate buffer) overnight at 4°C. Embryos were subsequently dissected in 30% sucrose and embedded in gelatin solution (7.5% gelatin, 15% sucrose in PB) and snap-frozen in isopentane on dry ice. Transverse cryosections (thickness: 12 mm) were cut using a Leica CM1950 cryostat in OCT and placed on Superfrost Plus slides. Slides were stored at -80C until ready to be processed for immunohistochemistry.

Slides were thawed at room temperature for 15mins before two immersions into PBS at 42°C to remove the gelatin. Slides were washed twice with PBST (PBS + 0.1% Triton-X100) for 5mins. Blocking was performed for 1hr with PBST + 2% BSA at room temperature, before incubation in PBST + 2% BSA with primary antibodies overnight at 4°C. Slides were washed 3 times for 5mins with PBST before incubation with secondary antibodies (see Table S1i) at room temperature for 1hr in the dark. After 3 PBST washes, incubation with Dapi (1/10 000) and washes with PBST and PBS, coverslips were mounted onto the sections using ProLong Gold antifade reagent and left overnight before imaging. Images were acquired on a Leica

Stellaris 5 or Leica SP8 STED confocal microscope. Primary antibodies are listed in Table S1b. Images presented in Figure 1C-F, 2I S1B,D are representative images of a minimum of three biological replicates. Images presented in S1F are representative of two biological replicates.

### Immunofluorescence on cells

Cells were washed in PBS and fixed in 4% paraformaldehyde in PBS for 30 mins at 4°C, followed by three washes in PBS. Primary antibodies (see Table S1b) were applied overnight at 4°C diluted in filtered blocking solution (2% BSA diluted in PBST – 0.1% Triton X-100 diluted in PBS). Cells were washed for 5 mins three times in PBST and incubated with secondary antibodies (see Table S1i) at room temperature, for 90 mins. Cells were washed for 5 mins three times in PBST, incubated with DAPI for 15 mins in PBS and washed twice before mounting with a glass coverslip using Prolong Gold (Invitrogen). Primary antibodies are listed in Table S1b.

Images were acquired on a Leica Stellaris 5 confocal microscope. Z stacks were acquired using the Leica LAS AF software and represented as maximum intensity projections using ImageJ software. The same settings were applied to all images. Images presented in Figure 3A, 4D, S3B are representative images of a minimum of three biological replicates.

### Flow cytometry

Cells were washed in PBS and dissociated with Accutase (Gibco). Once detached, cells were collected, washed with PBS, and pelleted. Cells were resuspended in PBS supplemented with live dye (1/1000, Thermo Fisher) and kept in dark at 4° for 30 mins. Cells were pelleted, washed in PBS, pelleted, and resuspended in 4% paraformaldehyde in PBS. Following 15 mins incubation at 4°C, cells were centrifuged, resuspended in PBS, and stored at 4°C for future analysis.

On the day of flow cytometry, cells were transferred for staining in U-bottom 96-well plates. Samples were pelleted and resuspended in 50μl block media (2% BSA diluted in PBST). After 30 mins incubation at room temperature in the platform rocker, antibodies were added to the sample and incubated overnight at 4°C on a platform rocker. Details of primary and secondary antibodies are described in Table S1c. Cells were pelleted for 4 mins, washed in PBST, pelleted, and incubated in 50μl PBST supplemented with secondary antibodies (concentration: 1/500) in the dark for 2h at room temperature in the platform rocker. One additional wash was performed before acquisition on a SymphonyA3 (BD Biosciences) using FACSDiva. Analysis was performed using FlowJo.

### RNA extraction, cDNA synthesis, and RT-qPCR analysis

RNA used for real time quantitative PCR (RT-qPCR) was extracted from cells using a QIAGEN RNeasy kit in RLT buffer, following the manufacturer’s instructions. Extracts were digested with DNase I to eliminate genomic DNA.

First-strand cDNA synthesis was performed using Superscript III (Invitrogen) using random hexamers and was amplified using PowerUp SYBR-Green Mastermix (Applied Biosystems). RT-qPCR was performed using the Applied Biosystems QuantStudio Real Time PCR system and analysed with Applied Biosystems QuantStudio 12K Flex software. PCR primers were designed using the online PrimerBLAST design tool and validated (standard curve and melting curve) or taken from previously published papers. Primer sequences are detailed in Table S1d. Two technical replicates were obtained for each sample and averaged before normalization and statistical analysis. Relative expression values for each gene were calculated by normalization against β-actin, using the delta–delta CT method. RT-qPCR analysis was performed on samples obtained from a minimum of three independent experiments for every primer pair analysed.

### mRNA-seq library preparation

RNA was extracted as previously described and RNA integrity was quality-controlled using the Bioanalyzer kit (RIN>9.7), mRNAseq libraries were made from 200ng total RNA using the Watchmaker Genomics Ltd mRNA Library Prep and RNA Library Prep kits according to manufacturer’s instructions. Libraries were dual indexed using IDT xGen™ Stubby Adapter-UDI Primers. Library quality and quantity was verified using the Agilent 4200 Tapestation and Qubit fluorometer respectively and then pooled in an equimolar manner and run on an Illumina NextSeq2000 sequencer, generating a minimum of 25 million Paired End 60bp reads per sample.

### ATAC-seq

ATAC-seq was performed following methods previously described (Metzis et al., 2018; Delas et al., 2023). Briefly, adherent cells were treated with StemPro Accutase to obtain a single cell suspension. Cells were counted and resuspended to obtain 5x10^4^ cells per sample in ice-cold PBS. Cells were pelleted and resuspended in lysis buffer (10mM Tris-HCl pH 7.4, 10mM NaCl, 3mM MgCl2, 0.1% IGEPAL, 0.1% Tween-20, 0.01% Digitonin) for 3mins on ice. Following a 10mins centrifugation at 4°C, nucleic extracts were resuspended in transposition buffer (supplemented with 0.1% Tween-20 and 0.01% digitonin) for 30mins at 37°C and purified using the QIAGEN MinElute PCR Purification kit following manufacturer’s instructions. Transposed DNA was eluted in a 20uL volume and amplified by PCR with Nextera indexes (see Table S1g) to generate single-indexed libraries. A maximum of 12 cycles of PCR was used to prevent saturation biases based on optimization experiments performed using qPCR. Library quality control was carried out using the Bioanalyzer High-Sensitivity DNA analysis kit. Libraries were pooled and sequenced as paired-end 50bp reads, on the NextSeq2000 platform. For all conditions, two or three biological replicate samples were collected from independent experiments.

### Spatial transcriptomics

*WT* C57BL6/J pregnant females were ordered from Charles River. Embryos were dissected in cold PBS, fixed for 45mins to 1hr in 4% PFA on ice under gentle agitation, dehydrated in 30% sucrose overnight at 4°C, embedded in OCT, sectioned in 10µm slices, and mounted onto Xenium slides according to 10X Genomics workflow CG000764.

Sections were then thawed and rehydrated using the 10X Genomics Visium CytAssist Spatial Gene Expression for Fixed Frozen workflow CG000662 with a modified thaw time of 1 min at 37°C. De-crosslinking was then performed as per 10X Genomics Xenium In Situ for FFPE workflow CG000580. Sections then underwent Priming Hybridisation, RNAse Treatment and Polishing, Probe Hybridisation, Ligation, Amplification, Cell Segmentation staining, Autofluorescence Quenching and Nuclei Staining according to 10X Genomics workflow CG000760: for Probe Hybridisation, the 10x Genomics pre-designed Xenium Prime 5K Mouse Pan Tissue and Pathways Gene Expression Panel was applied to the Xenium slides and hybridised for 17 hours at 50°C.

The optional multimodal Cell Segmentation was performed. During this step, the Xenium Multi-Tissue Stain Mix was applied to the sections for 16-20hrs at 4°C. This stain mix contains membrane-targeting antibodies, cytoplasmic antibodies, a universal ribosomal RNA interior stain, and the nuclear stain DAPI. These complementary signals are used as inputs for automated cell segmentation, enabling boundary definition firstly based on membrane labelling when available, or interior and nuclear signals when membrane labelling is insufficient. Autofluorescence Quenching was then performed to diminish unwanted autofluorescence and enhance signal-to-noise and nuclei are re-stained with DAPI. Finally, the samples were loaded into the Xenium Analyzer (according to 10X Genomics Protocol CG000584) for image processing, barcode decoding, and transcript localization using Xenium onboard analysis v3.3.1.1 (see Table S7b).

### Spatial transcriptomics data processing

Xenium data were loaded onto R studio (v4.4.1) using the Seurat (v5.3.0) function LoadXenium(). Initial filtering was performed by removing cells with 0 counts (nCount_Xenium > 0) followed by quality assessment of tissue integrity by looking at the images of the antibody segmentation mix and cell segmentation accuracy (Xenium Explorer) to remove scratched and significantly folded sections. Each section was manually labelled using the Xenium Explorer (v4.1.1). The co-ordinates were then exported (see Table S8a), loaded on R studio (v4.4.1), and transferred in the metadata to be used for downstream analysis. 1 indicates the 1st section taken and n indicates the n^th^ section taken (see Table S8b). Data were normalised using the SCTransform method (Seurat v5.3.0), which performs variance-stabilising transformation and regression of technical noise. Principal component analysis (PCA) was performed using RunPCA function at the default Seurat parameters to reduce dimensionality. The number of principal components of each used for downstream analysis was determined based on elbow plot for PC variance. Cell–cell similarity graphs were constructed using FindNeighbors(), followed by unsupervised clustering with FindClusters(). The clustering parameter (incl. resolution, knn, and dims) for each dataset was explored across multiple values, and cluster stability was assessed using the clustree (v0.5.1) package (Zappia et al., 2018). Cluster identities were assigned based on canonical marker gene expression (Table S9). To compare NCCs across datasets, extracted NCC clusters from each dataset were merged prior to integration generating a v5 object with multiple layers. That was followed by normalisation and PCA as described above. Batch effects and dataset-specific variation were corrected with the IntegrateLayers() function using the reciprocal PCA (RPCA) method (Stuart et al., 2019). An additional filtering step was performed for downstream integrated NCC analyses by removing cells with less than 100 counts (nCount_Xenium => 100).

### Differential gene expression analysis

Differential gene expression analysis was performed using the Seurat functions FindMarkers() and FindAllMarkers(), to identify differentially expressed genes for pairwise comparisons between specific clusters. Due to the limited number of cells in the Xenium data, the parameters were set as follows: logFC_threshold = 0.25, and at least 10% of cells expressing a given gene (min.pct = 0.10). Statistical testing was performed using the default Wilcoxon rank-sum test.

### Competing module analysis and visualization

Competing-lineage gene programmes were analysed using the sensory and autonomic gene modules from bifurcation 1 and the mesenchymal gene module from bifurcation 2 described by Soldatov et al. (2019) (Table S10a). Raw Xenium counts were library-size normalised per cell to 10,000 counts and log-transformed. For each module, genes not detected in the Xenium dataset were excluded, and per-cell module scores were calculated as the mean log-normalised expression of the remaining module genes.

For three-way comparison of competing transcriptional programmes, early sensory, autonomic and mesenchymal module scores were selected (Table S10b). Non-finite and negative values were set to zero before visualisation. For each cell, the three scores were normalised by their summed value, generating compositional proportions for sensory, autonomic and mesenchymal programmes. These proportions represent relative programme contribution rather than absolute module magnitude.

To compare transcriptional states across anatomical regions, normalised sensory, autonomic and mesenchymal proportions were projected onto a ternary simplex using barycentric coordinates. Cartesian coordinates were calculated from the three normalised proportions, positioning each cell according to its relative transcriptional composition. The vertices of the simplex correspond to “pure” sensory, autonomic and mesenchymal states, whereas intermediate positions represent mixed programme states.

To relate competing transcriptional states to axial identity, Hox paralogue group (PG1–PG13) scores were calculated as the mean log-normalised expression of detected Hox genes belonging to each paralogue group. Each cell was assigned a maximum Hox paralogue group (Max PG), defined as the paralogue group with the highest mean expression. Max PG identity was projected onto the ternary space and visualised using a discrete legend with one colour box per PG.

Regional occupancy of the ternary landscape was summarised by calculating convex hulls enclosing cells from each anatomical region. Two-dimensional kernel density contours were overlaid to visualise the distribution of cells within each regional territory.

For spatial visualisation, the same normalised sensory, autonomic and mesenchymal proportions were mapped to colourblind-friendly cyan, magenta and gold colour anchors, respectively. Colours were generated by weighted mixing of the three anchor colours according to the per-cell normalised programme proportions. Thus, colour represents relative programme composition. Cells below the signal threshold were shown in lighter grey compared to background cell types (other tissues). Spatial plotting was performed using ImageDimPlot() in Seurat.

### scRNA-seq data pre-processing and analysis

To analyse publicly available scRNA-seq data of mouse neural crest cells (GSE129114; Soldatov et al., 2019, Table S3), a standard bulk RNA-seq processing pipeline was applied, as unique molecular identifiers (UMIs) were not available. Quality control did not necessitate any corrective actions beyond confirming the use of Nextera adapter sequences, as indicated in the adapter content profiles. Raw sequencing reads were subjected to adapter trimming (Nextera adapters) and quality filtering (Phred score ≥ Q20) using Cutadapt (v4.7). The resulting high-quality reads were then quantified against the GRCm39 mouse transcriptome, using Ensembl release 110 annotations, via Salmon (v1.10.2). Gene expression matrices were filtered to remove low-quality cells using (nFeature_Count) thresholds as applied in Soldatov et al., 2019, where only cells with 3000 and 4000 total number of genes were kept, for E9.5 vagal+trunk *Wnt1* and E9.5 vagal+trunk *Sox10* datasets, respectively. Feature names were standardised by mapping Ensembl identifiers to gene symbols using org.Mm.eg.db with a biomaRt fallback. Features lacking valid gene symbols were removed, and duplicated gene symbols were collapsed by summing counts across corresponding entries. Normalisation and variable feature selection were performed independently for each subset using Seurat. Data were log-normalized (scale factor = 10,000), and the top 4000 variable features per plate were identified using the variance-stabilizing transformation (vst) method. Integration features were selected for all datasets, after which each of them was scaled using ScaleData() and subjected to PCA analysis as stated above. Data were integrated using Seurat’s RPCA method, with anchors identified using the first 50 principal components. The integrated assay was then scaled and re-analysed by PCA for downstream dimensionality reduction and clustering as previously described for the spatial transcriptomics.

### 5k Xenium probe panel testing in scRNAseq

To test whether the Xenium 5K gene panel can recapitulate full-transcriptome-defined NCC states, genes that were not included in the 5k panel were removed, the analysis was repeated as stated above, and concordance between annotated cell states and unsupervised clustering results was quantified. Cells were assigned cluster identities and overlap between reference cell states and Xenium-derived clusters was quantified. Values were normalised per cluster to show the proportion of cells assigned to each state. Quantitative agreement between annotations was assessed using the adjusted Rand index (ARI), providing a global measure of similarity between clustering assignments. Cluster purity was calculated as the proportion of cells belonging to the dominant reference state within each cluster. To further interpret cluster identities, each cluster was assigned to its predominant reference state based on majority scoring. These projected state labels were then mapped back to individual cells and visualised using UMAP embeddings.

### ATAC-seq analysis

Data was processed using the nf-core atacseq pipeline (https://nf-co.re/atacseq, Ewels et al., 2020) with the following options: –genome mm10 –macs_fdr 0.00001 -r 1.2.0 to generate bam and bigwig files.

For downstream analysis, fragments over 100bp were filtered out using samtools and peaks were called using genrich in ATAC-seq mode (https://github.com/jsh58/Genrich) and merged across replicates. After variance stabilizing transformation using DESeq2 (Love et al., 2014), replicates were compared using sample-to-sample Pearson correlation heatmap, principal component analysis and pair-wise comparison. Principal Component Analysis was performed using the top 10000 most variable elements and coloured by different sample metadata. Selection of differentially accessible elements across conditions was performed by pairwise DESeq2 analysis between any two cell types. Elements that fulfilled padj < 0.01 & abs(log2FoldChange) > 2 & baseMean > 100 were selected for subsequent clustering. Variance stabilized transformed data generated using DESeq2 were used as input to identify clusters of elements with the same dynamics. Clustering was performed using k-means with a high number of centers, and subsequently re-grouping clusters of very similar dynamics using hclust and target of 5 final clusters. Independent iterations resulted in reproducible clusters and dynamics. Using macs as an alternative peak caller resulted in similar clusters. Deeptools (Ramírez et al., 2016) was then used to map the coverage of ATAC-seq across these elements for the single and merged replicates. Homer was used for motif analysis across the differentially accessible elements using the findMotifsGenome function (using 200 as size of the region for motif finding, Heinz et al., 2010).

### mRNA-seq processing

mRNA-seq reads were processed using Cutadapt v4.7 (Martin, 2011) to remove adapter sequences and low-quality bases (minimum read length 31 bp; Phred score ≥20). Filtered reads were quantified with Salmon v1.10.2 (Patro et al., 2017) against the Ensembl GRCm39 mouse transcriptome (release 115), incorporating both cDNA and ncRNA annotations, with automatic library-type detection, soft clipping, and sequence- and GC-bias correction enabled. Transcript-level abundance estimates from 14 libraries were imported into R using tximport v1.38.2 and summarised to gene-level counts based on Ensembl gene identifiers derived from a transcript database generated with txdbmaker v1.6.2. Transcript version suffixes were removed prior to gene assignment. Genes with zero counts across all samples were excluded before downstream analyses.

Sample relationships and data quality were assessed using regularised log-transformed counts generated with DESeq2 v1.50.2. Principal component analysis (PCA) was performed using PCAtools v2.22.4.

Differential expression analysis was performed with DESeq2 v1.50.2 using a group-only design formula (∼0 + Group), treating biological replicates within each neural crest population as independent observations. Following library-size normalisation by size-factor estimation, counts were modelled using a negative binomial generalised linear model. Pairwise group comparisons were evaluated using Wald tests, and independent hypothesis weighting (IHW v1.38.0, Ignatiadis et al., 2016) was applied to improve statistical power. P values were adjusted for multiple testing using the Benjamini–Hochberg procedure, and log₂ fold-change estimates were shrunken using ashr (Stephens, 2017) for reporting and visualisation. No minimum fold-change threshold was applied during hypothesis testing. Genes with an adjusted P value < 0.01 were considered differentially expressed.

Functional enrichment analyses were conducted using the enricher function from clusterProfiler v4.18.4 on upregulated and downregulated gene sets from each contrast, defined by positive and negative log₂ fold changes, respectively. Gene sets were obtained from the mouse MSigDB collections via msigdbr v26.1.0 and included the Hallmark, Curated, Regulatory Target, Ontology, Immune Signature, and Cell Signature categories. Enrichment was assessed relative to the background set of all genes retained for differential expression testing. Statistical significance was evaluated using Benjamini–Hochberg correction, and gene sets with a nominal P value < 0.01 and a false discovery rate (q value) < 0.05 were considered significantly enriched.

---

**Figure S1:**
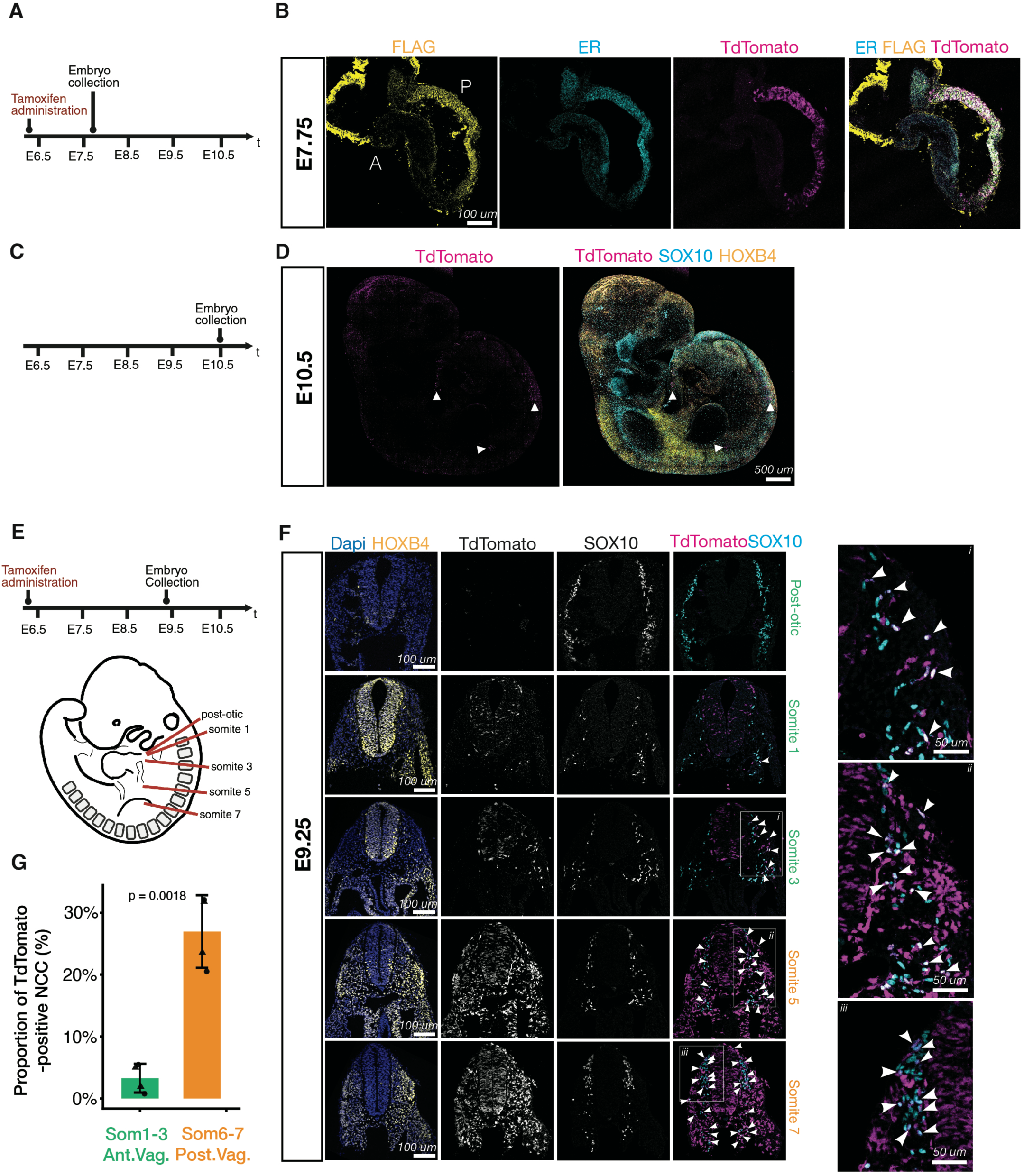
Validation of an inducible mouse line to track a *Cdx1* lineage. (A) Diagram of experimental approach to validate the lineage tracing mouse line. (B) Transverse sections of an E7.75 embryo shows co-localization of ER (cyan), FLAG (yellow) and TdTomato (magenta) in the posterior embryo (labelled P). (C) Experimental strategy to test Cre fidelity in the absence of tamoxifen administration. (D) WMIF of an E10.5 embryo shows TdTomato-positive (magenta) cells found at trunk and tail levels (white arrow) and not in cervical regions marked with HOXB4 (yellow). (E) Diagram of the experimental approach and schematic illustrating the distinct axial levels presented in panel F. (F) Transverse sections and immunofluorescence at E9.25 at indicated axial levels showing DAPI (blue), HOXB4 (yellow), TdTomato (magenta), and SOX10 (cyan). White boxes show magnified examples of co-localization (white arrows). (G) Quantification of TdTomato positive NCCs (labelled by SOX10) at the two most anterior and two most posterior vagal levels (n=2; t-test). ER, oestrogen receptor; WMIF, whole-mount immunofluorescence; E, embryonic day; NCCs, neural crest cells; DRG, dorsal root ganglia; SA, sympatho-adrenal; SE, sympatho-enteric. All schematic were created with BioRender.com.

**Figure S2:**
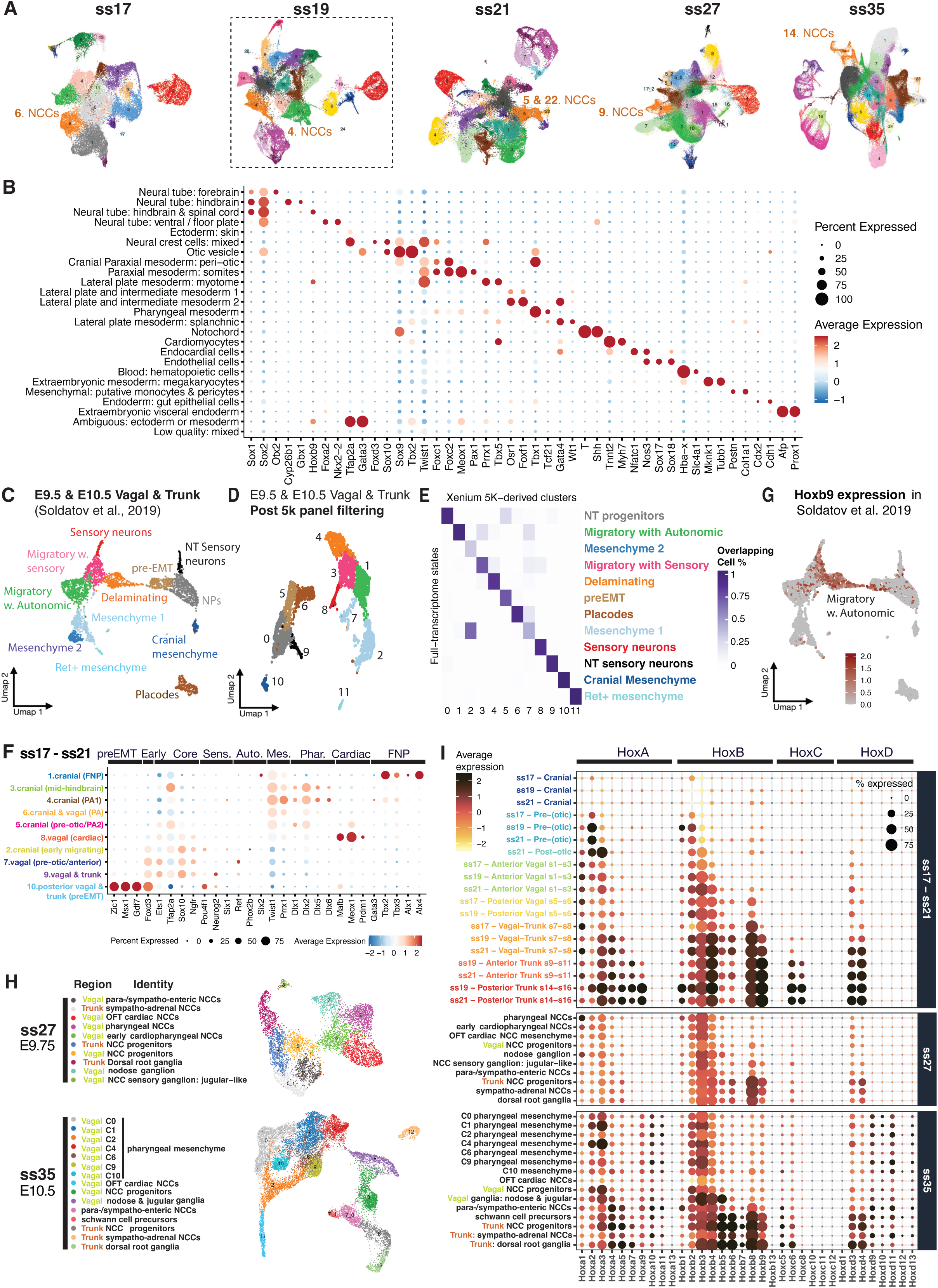
Identity profiling of neural crest using the spatial transcriptomics dataset. (A) UMAPs of each stage dataset coloured by cluster identity with highlighted putative NCC clusters. For each stage labels, see supplementary Tables S7a and S9. (B) Dot plot showing the expression of cell type-specific markers across all clusters extracted in the ss19 dataset (see A dashed lines). Dot size indicates the proportion of cells that express each gene and scale shows normalised gene expression. (C-D) UMAPs showing unsupervised clusters of extracted E9.5 and E10.5 NCCs at vagal and trunk level (Soldatov et al., 2019) using the full genome (C) or the Xenium 5k gene panel (D). (E) Heatmap showing state–cluster overlap comparing clusters derived from the 5k gene panel with full-transcriptome NCC states. Scale represents the proportion of cells that overlap across both datasets. (F) Heatmap of extracted NCCs between ss17 and 21 showing the expression of key NCC subtype markers obtained by Soldatov et al., 2019 and De Bono et al., 2021. (G) UMAP showing expression of Hoxb9 in extracted E9.5 and E10.5 NCCs at vagal and trunk level (Soldatov et al., 2019). (H) Annotated UMAPs of extracted NCC-like clusters for differentiating vagal and trunk derivatives (levels shown in Fig.2A) for ss27 and ss35. (I) Dot plot showing the expression of Hox genes across NCCs extracted from indicated time points and axial levels. Dot size indicates the proportion of cells that express each gene and scale shows normalised gene expression. UMAP, Uniform Manifold Approximation and Projection; ss, somite stage; s, somite; NCC(s), neural crest cell(s); E, embryonic day; SE, sympatho-enteric; SA, sympatho-adrenal.

**Figure S3:**
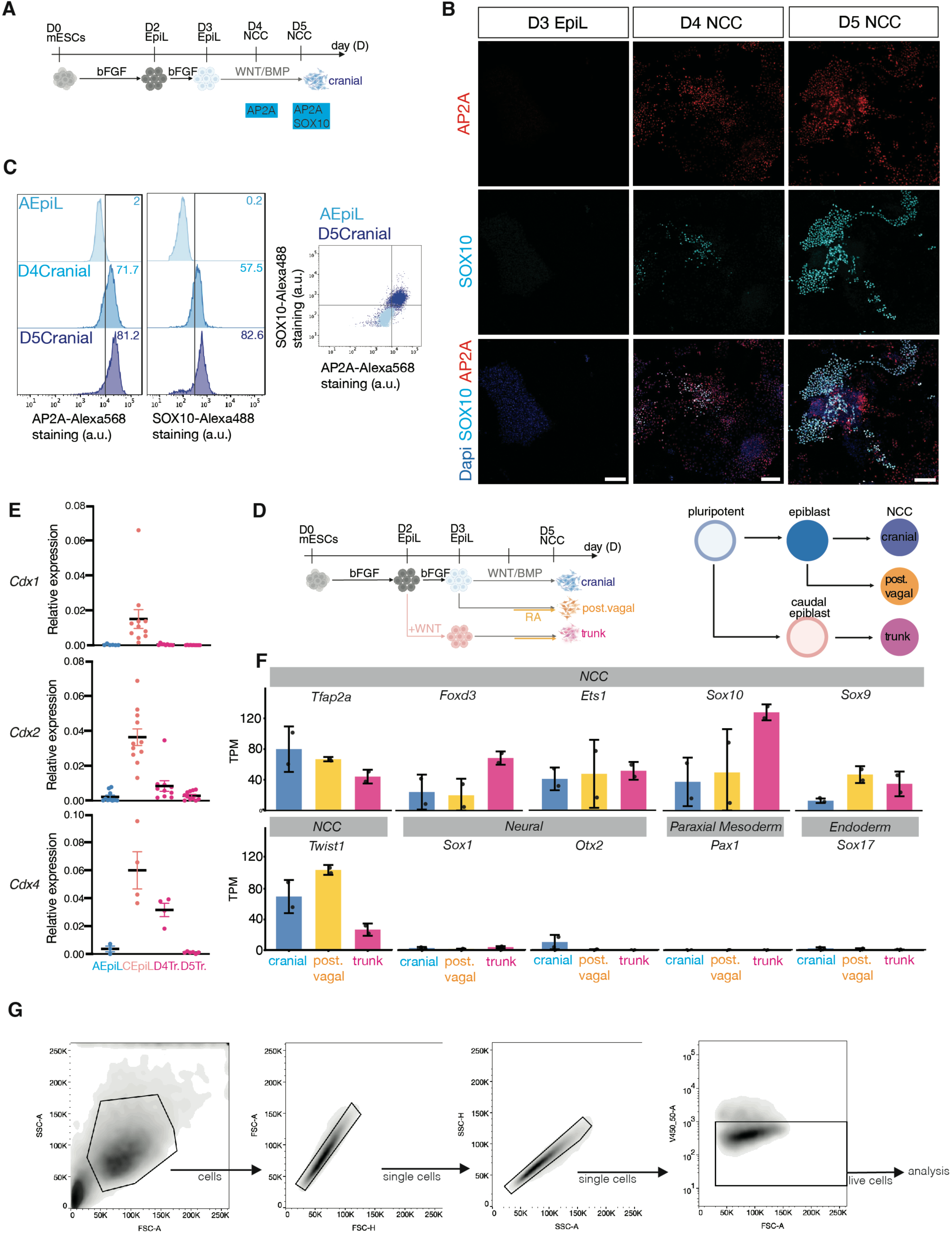
*In vitro* generation of neural crest cells. (A) Schematic of the 5-day cranial NCC differentiation from mESCs. (B-C) IF analysis (B) and cytometry histogram for TFAP2A and SOX10 with plot (C) of D4 and D5-NCC generated from EpiL shows progressive expression of TFAP2A by D4 (red) and SOX10 (cyan) by D5. (D) Schematic of the 5-day NCC differentiation from mESCs. (E) Relative expression (RT-qPCR) for *Cdx* genes shows transient expression of *Cdx1/2/4* as cells differentiate into trunk NCC progenitors. (F) Normalized expression levels assessed by mRNA-seq show that NCC generated *in vitro* express key NCC genes. (G) Gating strategy used for flow cytometry experiments. All schematic were created with BioRender.com. Data are represented as mean ± SEM. NCC, neural crest cell; mESCs, mouse embryonic stem cells; EpiL, epiblast-like; IF, immunofluorescence; D, day.

**Figure S4:**
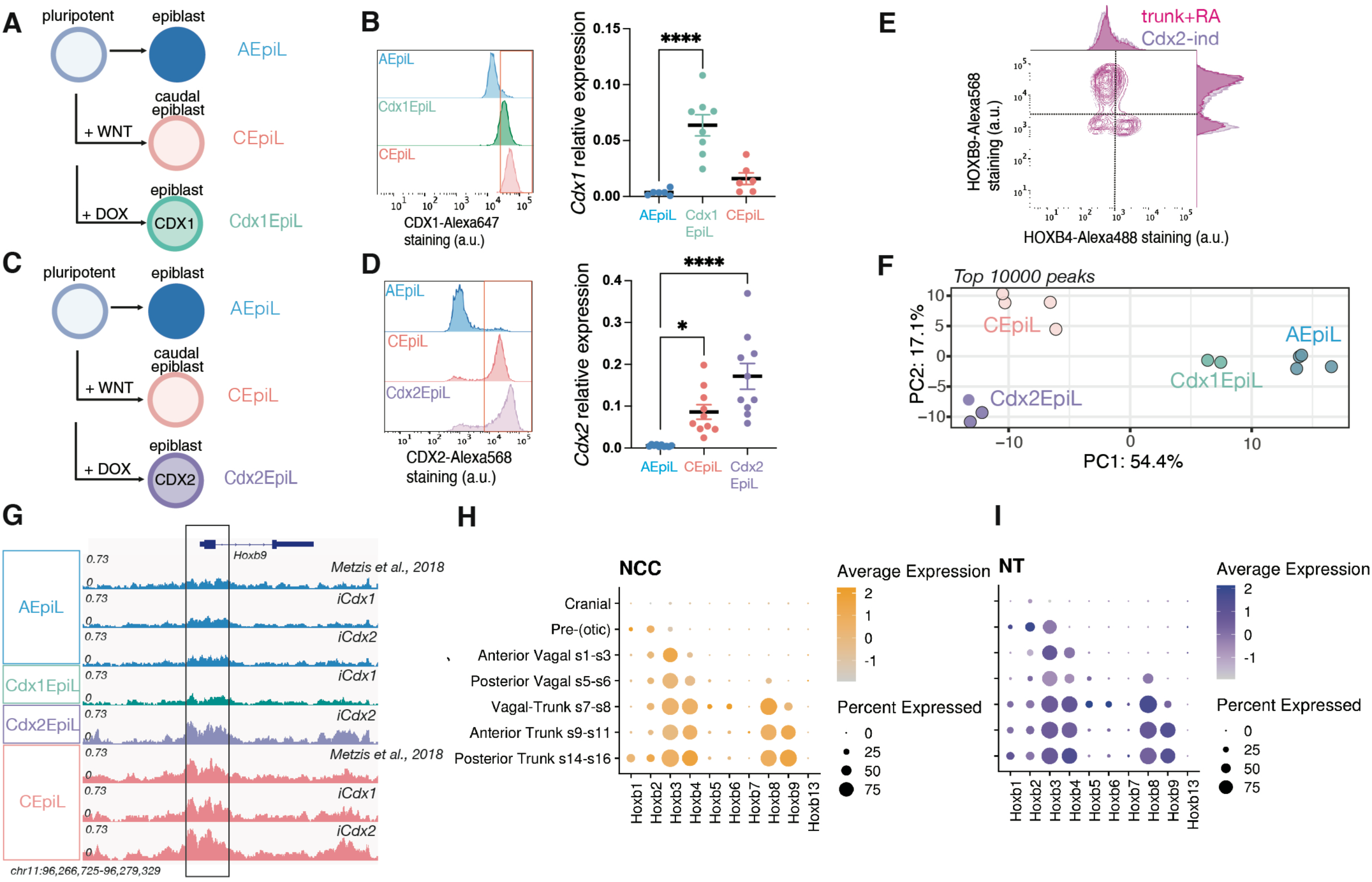
Differential induction of *Cdx1* and *Cdx2* changes chromatin accessibility. (A) Schematic of the 3-day EpiL differentiation from mESCs, (B) Cytometry histogram for CDX1 and *Cdx1* relative expression (RT-qPCR) shows upregulation of *Cdx1* in Cdx1EpiL generated from iCdx1 cells. (C) Schematic of the 3-day EpiL differentiation from mESCs. (D) Cytometry histogram for CDX2 and *Cdx2* relative expression (RT-qPCR) shows *Cdx2* upregulation in Cdx2EpiL generated from iCdx2 cells. (E) Representative flow cytometry plots showing TFAP2A-gated NCCs generated from *iCdx2* cells express only HOXB9 (Cdx2-ind and Trunk NCCs). (F) Dimensionality reduction (principal component analysis) based on the top 10,000 peaks shows induction of Cdx1 and Cdx2 differentially impacts chromatin accessibility: Cdx1EpiL (green) are closer to AEpiL (blue) while Cdx2 EpiL (purple) resemble CEpiL (pink). (G) Example locus that is becoming accessible in single Cdx2 (Cdx2EpiL) or combined Cdx (CEpiL) induction. (H-I) Dot plot of NCCs (G) and NPs (H) extracted from ss17-ss21 embryos grouped by regional identity showing normalised expression of the Hoxb genes. All schematic were created with BioRender.com. Data are represented as mean ± SEM. *p value < 0.05, **p value < 0.01, ****p value < 0.0001. EpiL, epiblast-like; mESCs, mouse embryonic stem cells; NCC, neural crest cell; CEpiL, caudal epiblast-like; AEpiL, anterior epiblast-like; NP, neural progenitors; SC, spinal cord; Hb, hindbrain.

## Notes

### Competing Interest Statement

The authors have declared no competing interest.

## References

1. Amblard, I. et al. A dual enhancer-attenuator element ensures transient Cdx2 expression during mouse posterior body formation. Developmental Cell 60, 2407–2419.e6 (2025).

2. Amin, S. et al. Cdx and T Brachyury Co-activate Growth Signaling in the Embryonic Axial Progenitor Niche. Cell Reports 17, 3165–3177 (2016).

3. Beck, F., Erler, T., Russell, A. & James, R. Expression of Cdx-2 in the mouse embryo and placenta: Possible role in patterning of the extra-embryonic membranes. Dev. Dyn. 204, 219–227 (1995).

4. Bel-Vialar, S., Itasaki, N. & Krumlauf, R. Initiating Hox gene expression: in the early chick neural tube differential sensitivity to FGF and RA signaling subdivides the *HoxB* genes in two distinct groups. Development 129, 5103–5115 (2002),

5. Blassberg, R. et al. Sox2 levels regulate the chromatin occupancy of WNT mediators in epiblast progenitors responsible for vertebrate body formation. Nat Cell Biol 24, 633–644 (2022).

6. Bolande, R. P. Neurocristopathy: Its Growth and Development in 20 Years. Pediatric Pathology & Laboratory Medicine 17, 1–25 (1997).

7. Bronner, M. E. & Simões-Costa, M. The Neural Crest Migrating into the Twenty-First Century. in Current Topics in Developmental Biology vol. 116 115–134.

8. Bronner-Fraser, M. & Fraser, S. E. Cell lineage analysis reveals multipotency of some avian neural crest cells. Nature 335, 161–164 (1988).

9. Calon, A. et al. Different effects of the Cdx1 and Cdx2 homeobox genes in a murine model of intestinal inflammation. Gut 56, 1688–1695 (2007).

10. Chan, W. Y., Cheung, C. S., Yung, K. M. & Copp, A. J. Cardiac neural crest of the mouse embryo: axial level of origin, migratory pathway and cell autonomy of the *splotch* (*Sp2H*) mutant effect. Development 131, 3367–3379 (2004).

11. Cheng, W. W.-C. et al. Depletion of the *IKBKAP* ortholog in zebrafish leads to hirschsprung disease-like phenotype. WJG 21, 2040–2046 (2015).

12. Conway, S. J. & Kaartinen, V. TGFβ superfamily signaling in the neural crest lineage. Cell Adhesion & Migration 5, 232–236 (2011).

13. Croaker, G. D. H., Shi, E., Simpson, E., Cartmill, T. & Cass, D. T. Congenital central hypoventilation syndrome and Hirschsprung’s disease. Archives of Disease in Childhood 78, 316–322 (1998).

14. Delás, M. J. et al. Developmental cell fate choice in neural tube progenitors employs two distinct cis-regulatory strategies. Developmental Cell 58, 3–17.e8 (2023).

15. Doetschman, T. et al. Targetted correction of a mutant HPRT gene in mouse embryonic stem cells. Nature 330, 576–578 (1987).

16. Domon-Dell, C. & Freund, J.-N. Stimulation of Cdx1 by oncogenic β-catenin/Tcf4 in colon cancer cells; opposite effect of the CDX2 homeoprotein. FEBS Letters 518, 83–87 (2002).

17. Duboule, D. & Dollé, P. The structural and functional organization of the murine HOX gene family resembles that of Drosophila homeotic genes. EMBO J 8, 1497–1505 (1989).

18. Durbec, P. L., Larsson-Blomberg, L. B., Schuchardt, A., Costantini, F. & Pachnis, V. Common origin and developmental dependence on *c-ret* of subsets of enteric and sympathetic neuroblasts. Development 122, 349–358 (1996).

19. 19. Erickson, A. G. et al. Unbiased profiling of multipotency landscapes reveals spatial modulators of clonal fate biases. Preprint at 10.1101/2024.11.15.623687 (2024).

20. Espinosa-Medina, I. et al. Dual origin of enteric neurons in vagal Schwann cell precursors and the sympathetic neural crest. Proc. Natl. Acad. Sci. U.S.A. 114, 11980–11985 (2017).

21. Ewels, Philip A et al. The nf-core framework for community-curatedbioinformatics pipelines. Nat Biotechnol 38, 271–278 (2020).

22. Fan, Y. et al. hPSC-derived sacral neural crest enables rescue in a severe model of Hirschsprung’s disease. Cell Stem Cell 30, 264–282.e9 (2023).

23. Fattahi, F. et al. Deriving human ENS lineages for cell therapy and drug discovery in Hirschsprung disease. Nature 531, 105–109 (2016).

24. Frith, T. J. R. et al. Retinoic Acid Accelerates the Specification of Enteric Neural Progenitors from In-Vitro-Derived Neural Crest. Stem Cell Reports 15, 557–565 (2020).

25. Frith, T. J. et al. Human axial progenitors generate trunk neural crest cells in vitro. eLife 7, e35786 (2018).

26. Gaultier, C., Trang, H., Dauger, S. & Gallego, J. Pediatric Disorders with Autonomic Dysfunction: What Role for PHOX2B? Pediatr Res 58, 1–6 (2005).

27. Gaunt, S. J., Drage, D. & Trubshaw, R. C. cdx4/lacZ and cdx2/lacZ protein gradients formed by decay during gastrulation in the mouse. Int. J. Dev. Biol. 49, 901–908 (2005).

28. Gogolou, A. et al. Early anteroposterior regionalisation of human neural crest is shaped by a pro-mesodermal factor. eLife 11, e74263 (2022).

29. Gouti, M. et al. In Vitro Generation of Neuromesodermal Progenitors Reveals Distinct Roles for Wnt Signalling in the Specification of Spinal Cord and Paraxial Mesoderm Identity. PLoS Biol 12, e1001937 (2014).

30. Grainger, S., Hryniuk, A. & Lohnes, D. Cdx1 and Cdx2 Exhibit Transcriptional Specificity in the Intestine. PLoS ONE 8, e54757 (2013).

31. Green, S. A. Ancient evolutionary origin of vertebrate enteric neurons from trunk-derived neural crest.

32. 32. Hackland, J. O. S. et al. Top-Down Inhibition of BMP Signaling Enables Robust Induction of hPSCs Into Neural Crest in Fully Defined, Xeno-free Conditions. Stem Cell Reports **9**, 1043–1052 (2017).

33. Heinz, S. et al. Simple Combinations of Lineage-Determining Transcription Factors Prime cis-Regulatory Elements Required for Macrophage and B Cell Identities. Molecular Cell 38, 576–589 (2010).

34. Howard, A. G. A. et al. Elevated Hoxb5b Expands Vagal Neural Crest Pool and Blocks Enteric Neuronal Development in Zebrafish. Front. Cell Dev. Biol. 9, 803370 (2022).

35. Howard, A. G. et al. An atlas of neural crest lineages along the posterior developing zebrafish at single-cell resolution. eLife 10, e60005 (2021).

36. Hsu, I.-U. Y. et al. Lineage and organ signals sequentially build organ intrinsic nervous systems. Nature 655, 429–437 (2026).

37. Hutchins, E. J. et al. Migration and diversification of the vagal neural crest. Developmental Biology 444, S98–S109 (2018).

38. Ignatiadis, N., Klaus, B., Zaugg, J. B. & Huber, W. Data-driven hypothesis weighting increases detection power in genome-scale multiple testing. Nat Methods 13, 577–580 (2016).

39. Ishikawa, S. & Ito, K. Plasticity and regulatory mechanisms of Hox gene expression in mouse neural crest cells. Cell Tissue Res 337, 381–391 (2009).

40. Kirby, Margaret L., Gale, Thomas F., & Stewart, Donald E. Neural Crest Cells Contribute to Normal Aorticopulmonary Septation. Science 220, 1059–1061 (1983).

41. Krispin, S., Nitzan, E., Kassem, Y. & Kalcheim, C. Evidence for a dynamic spatiotemporal fate map and early fate restrictions of premigratory avian neural crest. Development 137, 585–595 (2010).

42. Kuo, B. R. & Erickson, C. A. Vagal neural crest cell migratory behavior: A transition between the cranial and trunk crest. Developmental Dynamics 240, 2084–2100 (2011).

43. Le Douarin, N. M. & Teillet, M.-A. M. Experimental analysis of the migration and differentiation of neuroblasts of the autonomic nervous system and of neurectodermal mesenchymal derivatives, using a biological cell marking technique. Developmental Biology 41, 162–184 (1974).

44. Le Douarin NM, Teillet MA. The migration of neural crest cells to the wall of the digestive tract in avian embryo. J Embryol Exp Morphol. 30, 31–48 (1973).

45. Ling, I. T. C. & Sauka-Spengler, T. Early chromatin shaping predetermines multipotent vagal neural crest into neural, neuronal and mesenchymal lineages. Nat Cell Biol 21, 1504–1517 (2019).

46. Lippmann, E. S. et al. Deterministic HOX Patterning in Human Pluripotent Stem Cell-Derived Neuroectoderm. Stem Cell Reports 4, 632–644 (2015).

47. Love, M. I., Huber, W. & Anders, S. Moderated estimation of fold change and dispersion for RNA-seq data with DESeq2. Genome Biol 15, 550 (2014).

48. Lukoseviciute, M., Mayes, S. & Sauka-Spengler, T. Neuromesodermal progenitor origin of trunk neural crest in vivo.

49. Ma, Q., Kintner, C. & Anderson, D. J. Identification of neurogenin, a Vertebrate Neuronal Determination Gene. Cell 87, 43–52 (1996).

50. Madisen, L. et al. A robust and high-throughput Cre reporting and characterization system for the whole mouse brain. Nat Neurosci 13, 133–140 (2010).

51. Mallo, M., Wellik, D. M. & Deschamps, J. Hox genes and regional patterning of the vertebrate body plan. Developmental Biology 344, 7–15 (2010).

52. Marcel Martin. Cutadapt removes adapter sequences from high-throughoutputs sequencing reads. (2011).

53. Mariani, L. et al. A TALE/HOX code unlocks WNT signalling response towards paraxial mesoderm. Nat Commun 12, 5136 (2021).

54. Metzis, V. et al. Nervous System Regionalization Entails Axial Allocation before Neural Differentiation. Cell 175, 1105–1118.e17 (2018).

55. Meyer, B. I. & Gruss, P. Mouse *Cdx-1* expression during gastrulation. Development 117, 191–203 (1993).

56. Mouilleau, V. et al. Dynamic extrinsic pacing of the HOX clock in human axial progenitors controls motor neuron subtype specification. Development 148, dev194514 (2021).

57. Nagashimada, M. et al. Autonomic neurocristopathy-associated mutations in PHOX2B dysregulate Sox10 expression. J. Clin. Invest. 122, 3145–3158 (2012).

58. Needham, J. & Metzis, V. Heads or tails: making the spinal cord. Developmental Biology 485, 80–92 (2022).

59. Parker, H. J., Bronner, M. E. & Krumlauf, R. The vertebrate *Hox* gene regulatory network for hindbrain segmentation: Evolution and diversification: Coupling of a *Hox* gene regulatory network to hindbrain segmentation is an ancient trait originating at the base of vertebrates. BioEssays 38, 526–538 (2016).

60. Patro, R., Duggal, G., Love, M. I., Irizarry, R. A. & Kingsford, C. Salmon provides fast and bias-aware quantification of transcript expression. Nat Methods 14, 417–419 (2017).

61. Ramírez, F. et al. deepTools2: a next generation web server for deep-sequencing data analysis. Nucleic Acids Res 44, W160–W165 (2016).

62. Rocha, M. et al. Zebrafish Cdx4 regulates neural crest cell specification and migratory behaviors in the posterior body. Developmental Biology 480, 25–38 (2021).

63. Sanchez-Ferras, O. et al. A direct role for murine Cdx proteins in the trunk neural crest-gene regulatory network. Development dev.132159 (2016) doi:10.1242/dev.132159.

64. Savory, J. G. A. et al. Cdx1 and Cdx2 are functionally equivalent in vertebral patterning. Developmental Biology 330, 114–122 (2009).

65. Serafimidis, I., Rakatzi, I., Episkopou, V., Gouti, M. & Gavalas, A. Novel Effectors of Directed and Ngn3-Mediated Differentiation of Mouse Embryonic Stem Cells into Endocrine Pancreas Progenitors. Stem Cells 26, 3–16 (2008).

66. Simkin, J. E., Zhang, D., Stamp, L. A. & Newgreen, D. F. Fine scale differences within the vagal neural crest for enteric nervous system formation. Developmental Biology 446, 22–33 (2019).

67. Simoes-Costa, M. & Bronner, M. E. Reprogramming of avian neural crest axial identity and cell fate. Science 352, 1570–1573 (2016).

68. Simon, C. S. & Hadjantonakis, A.-K. Top to Tail: Anterior-Posterior Patterning Precedes Regional Nervous System Identity. Cell 175, 905–907 (2018).

69. Soldatov, R. et al. Spatiotemporal structure of cell fate decisions in murine neural crest. Science 364, eaas9536 (2019).

70. Stephens, M. False discovery rates: a new deal. Biostatistics (2017) 18, 2, doi:10.1093/biostatistics/kxw041

71. Subramanian, V., Meyer, B. I. & Gruss, P. Disruption of the murine homeobox gene Cdx1 affects axial skeletal identities by altering the mesodermal expression domains of Hox genes. Cell 83, 641–653 (1995).

72. Tang, W., Li, Y., Li, A. & Bronner, M. E. Clonal analysis and dynamic imaging identify multipotency of individual Gallus gallus caudal hindbrain neural crest cells toward cardiac and enteric fates. Nat Commun 12, 1894 (2021).

73. Van Den Akker, E., et al. *Cdx1* and *Cdx2* have overlapping functions in anteroposterior patterning and posterior axis elongation. Development 129, 2181–2193 (2002).

74. Van Rooijen, C. et al. Evolutionarily conserved requirement of Cdx for post-occipital tissue emergence. Development 139, 2576–2583 (2012).

75. Vega-Lopez, G. A., Cerrizuela, S., Tribulo, C. & Aybar, M. J. Neurocristopathies: New insights 150 years after the neural crest discovery. Developmental Biology 444, S110–S143 (2018).

76. Vong, K. I. et al. Developmental organization of sensory and sympathetic ganglia. Nature 654, 734–743 (2026). 10.1038/s41586-026-10313-0

77. Young, T. et al. Cdx and Hox Genes Differentially Regulate Posterior Axial Growth in Mammalian Embryos. Developmental Cell 17, 516–526 (2009).

78. Zalc, A. et al. Reactivation of the pluripotency program precedes formation of the cranial neural crest. Science 371, eabb4776 (2021).

79. Ziwei Liu et al. Resolving cell lineages and gene functions in the developing mouse gastrointestinal tract using in utero transduction. Preprint 10.64898/2026.04.09.716777 (2026)

